# A Structure-Based Mechanism for DNA Entry into the Cohesin Ring

**DOI:** 10.1101/2020.04.21.052944

**Authors:** Torahiko L. Higashi, Patrik Eickhoff, Joana S. Simoes, Julia Locke, Andrea Nans, Helen R. Flynn, Ambrosius P. Snijders, George Papageorgiou, Nicola O’Reilly, Zhuo A. Chen, Francis J. O’Reilly, Juri Rappsilber, Alessandro Costa, Frank Uhlmann

**Affiliations:** Chromosome Segregation Laboratory; Macromolecular Machines Laboratory; Structural Biology Science Technology Platform; Proteomics Science Technology Platform; Peptide Chemistry Science Technology Platform, The Francis Crick Institute, London NW1 1AT, UK; Bioanalytics Unit, Institute of Biotechnology, Technische Universität Berlin, 13355 Berlin, Germany; Wellcome Centre for Cell Biology, University of Edinburgh, Edinburgh EH9 3BF, UK

**Keywords:** Chromosome segregation, Sister chromatid cohesion, SMC complexes, Cohesin, ABC-ATPase, Mis4/Scc2/NIPBL, Cryo-electron microscopy, DNA-protein crosslink mass spectrometry, *S*. *pombe*

## Abstract

Despite key roles in sister chromatid cohesion and chromosome organization, the mechanism by which cohesin rings are loaded onto DNA is still unknown. Here, we combine biophysical approaches and cryo-EM to visualize a cohesin loading intermediate in which DNA is locked between two gates that lead into the cohesin ring. Building on this structural framework, we design biochemical experiments to establish the order of events during cohesin loading. In an initial step, DNA traverses an N-terminal kleisin gate that is first opened upon ATP binding and then closed as the cohesin loader locks the DNA against a shut ATPase gate. ATP hydrolysis leads to ATPase gate opening to complete DNA entry. Whether DNA loading is successful, or rather results in loop extrusion, might be dictated by a conserved kleisin N-terminal tail that guides the DNA through the kleisin gate. Our results establish the molecular basis for cohesin loading onto DNA.

## INTRODUCTION

The SMC protein family is conserved from prokaryotic to eukaryotic cells and their role in DNA organization is vital for chromosome function. Eukaryotes contain three distinct SMC family members, cohesin, condensin and the Smc5-Smc6 complex (Hirano, 2016; Jeppsson et al., 2014; Uhlmann, 2016). All SMC complexes possess a similar large ring structure that is adapted to the overlapping, yet distinct and essential functions that each of the complexes fulfil. Among them, the cohesin complex establishes cohesion between replicated sister chromatids, which forms the basis for their faithful segregation during cell division. Additional roles of cohesin include chromatin domain organization in interphase, as well as DNA repair by homologous recombination. The overarching role of cohesin in these activities is thought to be the establishment of interactions between more than one DNA. A characteristic feature of cohesin is its ability to bind DNA by topological embrace, which underpins sister chromatid cohesion (Haering et al., 2008). At the same time, cohesin was seen to extrude DNA loops, without need for the ring to topologically trap DNA (Davidson et al., 2019; Kim et al., 2019). Such a loop extrusion mechanism has been proposed to underlie interphase chromatin domain organization. The molecular mechanism by which cohesin topologically entraps DNA, or extrudes a DNA loop, is not yet understood.

The cohesin ring consists of three core components, two SMCs and a kleisin subunit. The two SMC subunits, Psm1^Smc1^ and Psm3^Smc3^, form long anti-parallel coiled coils, which stably interact at one end at a dimerization motif, called the ‘hinge’. The SMC coiled coils show flexibility, pivoting at an ‘elbow’ that is situated approximately half-way along their length (Anderson et al., 2002; Bürmann et al., 2019). At the other end of the coiled coil lie ATP-binding cassette (ABC)-type nucleotide binding domains, known as ‘heads’. These heads dimerize following ATP binding and disengage upon ATP hydrolysis (Hopfner et al., 2000). Whether the cohesin ATPase is engaged or disengaged in its resting state is not yet known. Α kleisin subunit, Rad21^Scc1^, bridges the two ATPase heads to complete this ring architecture. Elements close to the kleisin N-terminus form a triple helix with the Psm3 ‘neck’ where the coiled coil joins the Psm3 ATPase head. The kleisin C-terminus in turn forms a small winged-helix domain that associates with the bottom of the Psm1 head (Gligoris et al., 2014; Huis in ‘t Veld et al., 2014). This asymmetric kleisin binding to the ATPase heads was first seen in prokaryotic SMC proteins (Bürmann et al., 2013), suggesting a conserved molecular architecture and common principles of SMC complex function.

The SMC-kleisin ring is regulated by three additional HEAT repeat subunits that associate with the unstructured middle region of the kleisin. The Psc3^Scc3/STAG1/2^ subunit binds to the center of this region and is instrumental for both cohesin loading and unloading (Hara et al., 2014; Li et al., 2018; Murayama and Uhlmann, 2014, 2015). Mis4^Scc2/NIPBL^ is known as the cohesin loader. It binds the kleisin upstream of Psc3^Scc3/STAG1/2^ and together with its Ssl3^Scc4/MAU2^ binding partner is essential for chromosomal cohesin loading. Mis4^Scc2/NIPBL^ by itself is sufficient to promote *in vitro* cohesin loading onto DNA, while Ssl3^Scc4/MAU2^ serves as an *in vivo* chromatin adaptor (Chao et al., 2015; Furuya et al., 1998; Kikuchi et al., 2016; Muñoz et al., 2019; Murayama and Uhlmann, 2014). The HEAT subunit Pds5, in turn, competes with Mis4^Scc2/NIPBL^ for kleisin binding. Pds5 has a dual role in both stabilizing loaded cohesin on DNA, as well as recruiting a binding partner, Wapl, that promotes cohesin unloading (Lee et al., 2016; Murayama and Uhlmann, 2015; Ouyang et al., 2016; Tanaka et al., 2001). Thus the loader and unloader pair influences cohesin in opposing ways, resulting in dynamic cohesin turnover on chromosomes. This changes during DNA replication when two conserved Psm3^Smc3^ lysine residues (K105-K106 in fission yeast) are acetylated by the replication fork-associated Eso1^Eco1/ESCO2^ acetyltransferase (Ben-Shahar et al., 2008; Ünal et al., 2008; Zhang et al., 2008). Acetylation stabilizes cohesin binding to chromosomes, which is a requirement for enduring sister chromatid cohesion.

Recent studies have started to shed light onto the molecular mechanism of cohesin unloading from DNA. Cohesin unloading depends on ATP hydrolysis and therefore likely on dissociation of the ATPase heads (head gate opening). Pds5 and Wapl in turn enact dissociation of the kleisin N-terminus from Psm3^Smc3^ (kleisin N-gate opening), consistent with an outward DNA trajectory through the ATPase head and kleisin N-gates (Beckouët et al., 2016; Buheitel and Stemmann, 2013; Chan et al., 2012; Murayama and Uhlmann, 2015). Psm3^Smc3^ K105-K106 mediate DNA stimulated ATP hydrolysis, offering a rationale for how their acetylation stabilizes cohesin on chromosomes. While head gate opening depends on ATP hydrolysis, kleisin N-gate opening requires head engagement (Murayama and Uhlmann, 2015).

We therefore proposed an “inter-locking gate” model in which DNA exits the SMC ring through sequential head and kleisin N-gates, only one of which can be open at any one time. The exact sequence of events that lead to cohesin unloading, however, remains to be ascertained.

How DNA enters the cohesin ring remains controversial. Just like unloading, cohesin loading onto DNA is coupled to its ATPase function and the Mis4^Scc2/NIPBL^ cohesin loader stimulates DNA-dependent ATP hydrolysis (Arumugam et al., 2003; Murayama and Uhlmann, 2014; Weitzer et al., 2003). Psm3 K105-K106 are also required and, at least *in vitro*, Pds5-Wapl likewise facilitate topological loading (Murayama and Uhlmann, 2015). As these requirements are similar to those of cohesin unloading, we hypothesized that cohesin loading uses the same DNA trajectory through the interlocking ATPase head and kleisin N-gates. The entry reaction would be facilitated by loader-dependent cohesin ring folding, exposing the luminal Psm3^Smc3^ K105-K106 residues to DNA (Murayama and Uhlmann, 2015).

However, *in vitro* studies suggest that ATP binding but not hydrolysis is required for topological cohesin binding to DNA, which is hard to reconcile with the above model (Çamdere et al., 2018; Minamino et al., 2018). Furthermore, fusing budding yeast Smc3 to the kleisin N-terminus in one polypeptide chain still allowed chromosome binding of cohesin (Gruber et al., 2006), which was interpreted to rule out DNA passage through the kleisin N-gate. Ligand-mediated crosslinking of hinge insertions in turn prevented cohesin loading, suggesting that DNA enters through the hinge (Buheitel and Stemmann, 2013; Gruber et al., 2006).

To understand the process of cohesin loading onto DNA, we used Förster resonance energy transfer (FRET) to measure conformational changes at the SMC heads. This showed that the SMC heads engage in the presence of DNA, the cohesin loader and non-hydrolyzable ATP. We visualized this cohesin loading intermediate by cryo-electron microscopy (EM) at an average resolution of 3.9 Å. DNA is trapped between the kleisin N-gate and the head gate, while the cohesin loader plays a key structural role in stabilizing this state. The development of DNA-protein crosslink mass spectrometry (DPC-MS) allows us to trace the DNA trajectory, leading to a model whereby DNA passes through the kleisin N-gate before reaching the engaged SMC heads. DNA and the loader will trigger ATP hydrolysis and head disengagement to complete DNA entry into the cohesin ring.

## RESULTS

### Cohesin ATPase Head Engagement with Loader, DNA and ATP

To understand how ATPase head engagement by cohesin is regulated, we monitored head proximity of purified fission yeast cohesin using FRET. We used a tetramer complex consisting of Psm1, Psm3, Rad21 and Psc3 (Murayama and Uhlmann, 2014) including C-terminal SNAP and CLIP-tags on Psm1 and Psm3, respectively, that could be specifically labelled with Dy547 and Alexa 647 fluorophores as a FRET pair (Figures 1A, S1A). The labelled cohesin displayed wild type levels of Mis4-dependent DNA loading and (Figure S1B). In the absence of ATP and DNA, cohesin exhibited measurable FRET, suggesting relative proximity between the two ATPase heads. Addition of ATP slightly increased FRET efficiency, consistent with ATP-dependent head engagement (Figure S1C). Combination of ATP with a 3 kb circular plasmid DNA and loader did not further augment the FRET signal, though we note that addition of the Mis4-Ssl3 cohesin loader alone or in pairwise combinations with ATP or DNA reproducibly reduced FRET efficiency.

**Figure 1.**
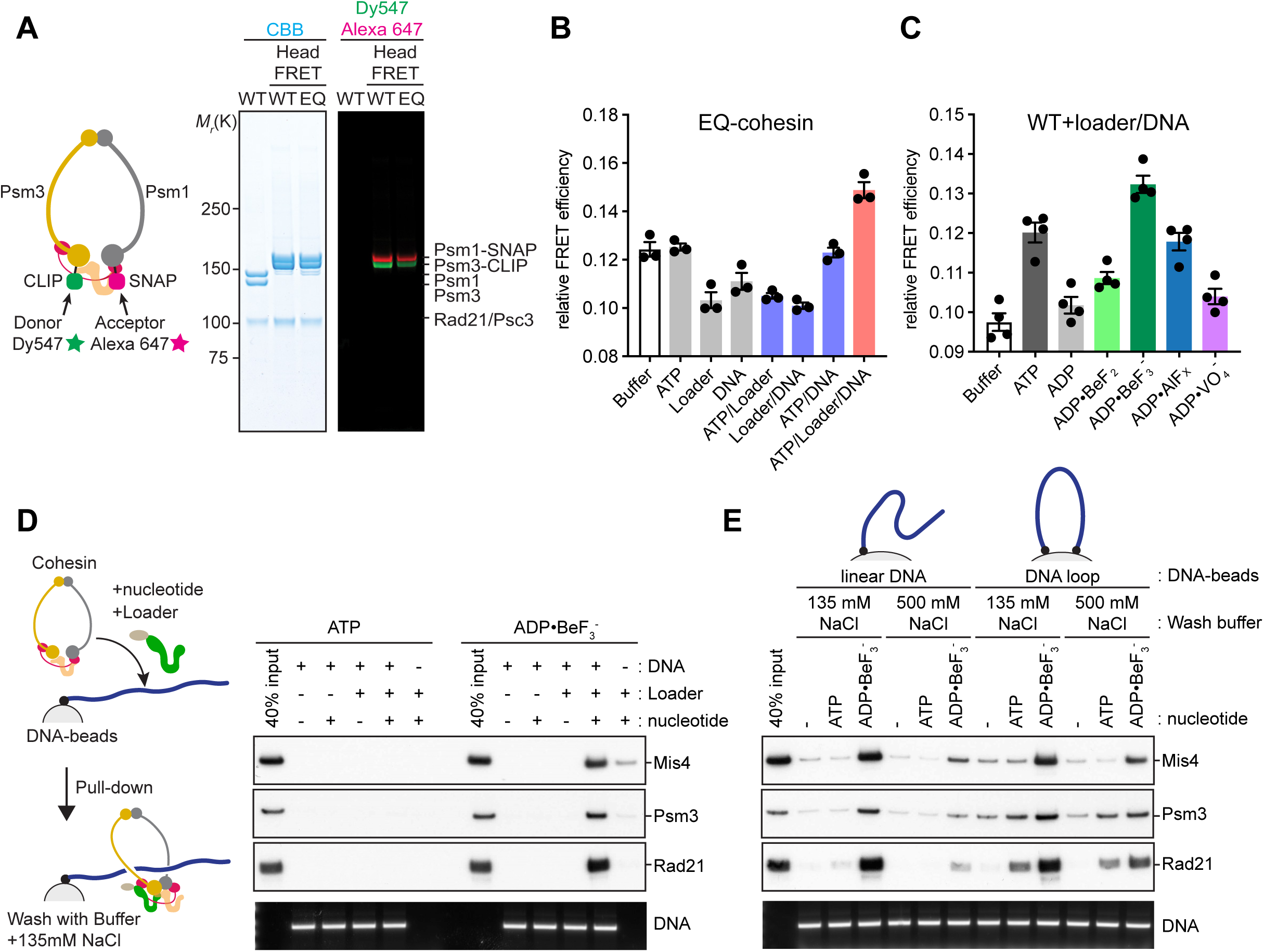
Cohesin ATPase Head Engagement Leads to a DNA ‘Gripping’ State (A) Schematic, purification and labeling of untagged, and fluorescent tagged wild type (WT) and Walker B mutant (EQ) cohesin to measure FRET between the Psm1 and Psm3 ATPase heads. Purified and labeled complexes were analyzed by SDS-polyacrylamide gel electrophoresis (SDS-PAGE) followed by Coomassie blue (CBB) staining or in gel fluorescence detection. (B) Head FRET efficiencies of EQ-cohesin with the indicated additions was calculated by dividing the Alexa 647 intensity at its emission peak with the sum of Alexa 647 and Dy547 intensities. Results from three independent repeats of the experiment, their means and standard deviations are shown. (C) Head FRET efficiencies of wild type cohesin in the presence of the Mis4-Ssl3 loader, a 3 kb circular plasmid DNA and the indicated nucleotides and phosphate analogs was measured as in (B). Results from four independent repeats of the experiment, their means and standard deviations are shown. (D) Schematic of the DNA gripping experiment in which biotinylated linear dsDNA is immobilized on streptavidin-coated magnetic beads. Cohesin, loader and the indicated nucleotides were added. After the incubation, beads were washed with buffer containing 135 mM NaCl. Bound protein was analyzed by SDS-PAGE and immunoblotting while the DNA was visualized by agarose gel electrophoresis. (E) Salt sensitivity of cohesin-DNA complexes following assembly with hydrolyzable or non-hydrolyzable ATP, on linear DNA and on DNA loops. Following the incubation, beads were washed with buffer containing 135 mM or 500 mM NaCl and products analyzed as in (D).

Cycles of head engagement and disengagement following ATP hydrolysis might dampen any bulk FRET changes. To prevent ATPase cycling, we purified ATP hydrolysis-deficient Walker B motif mutant cohesin (EQ-cohesin; (Murayama and Uhlmann, 2015); Figure 1A). FRET changes were indeed augmented when using EQ-cohesin. Addition of Mis4-Ssl3, with or without DNA, again resulted in FRET loss (Figure 1B), suggesting that the cohesin loader has a tendency to separate the ATPase heads. Strikingly, the presence of all three components, the loader, DNA and ATP, resulted in a marked FRET increase. This suggests that head engagement is reached when all loading components come together. We observed this state only using ATP hydrolysis-deficient cohesin, indicating that head engagement is usually transient. A further implication of this observation is that, at most times, the ATPase heads of wild type cohesin are in proximity but not engaged.

To explore the requirements for head engagement further, we compared the Mis4-Ssl3 cohesin loader complex with a truncation mutant of Mis4, lacking its N-terminal Ssl3 interacting region (Mis4-N191; (Chao et al., 2015)). Mis4-N191 is competent in *in vitro* cohesin loading and was equally proficient in promoting head engagement (Figure S1D). On the contrary, Pds5-Wapl did not promote head engagement, thus revealing a mechanistic difference between loader and unloader (Figure S1E). When studying the DNA requirement, we found linear or circular double stranded DNA (dsDNA) to have equal effects, while circular single stranded DNA (ssDNA) was slightly more efficient at promoting head engagement (Figure S1F). Thus the loader and either dsDNA or ssDNA cooperatively promote ATP-dependent head engagement.

In addition to EQ-cohesin, the wild type cohesin complex reaches topological DNA binding in the presence of ADP and phosphate analogs that mimic the ATP ground state (Çamdere et al., 2018; Minamino et al., 2018; Murayama and Uhlmann, 2015). To confirm that head engagement is similarly reached under these conditions, we measured FRET of wild type cohesin with the loader, DNA, ADP and *<ι>γ</i>*-phosphate analogs (Figure 1C). The ATP ground state mimics BeF_3_ and AlF_x_, but not BeF_2_ or the transition state mimic VO_4_, elicited a FRET increase, consistent with SMC head engagement following ATP binding in its ground state.

### Head Engagement Leads to a DNA ‘Gripping’ State

While performing the above experiments, we noticed that cohesin binds unusually tight to linear DNA in the presence of loader and non-hydrolyzable ATP. When using linear bead-bound DNA as a substrate for cohesin loading, little cohesin is retained following incubation with loader and ATP and a wash at physiological salt concentration (135 mM NaCl). However, when we used ADP·BeF_3_ as the nucleotide, both cohesin and the loader were efficiently retained on DNA (Figure 1D). Tight DNA binding was reproduced using ATP and EQ-cohesin (Figure S1G). Mis4-N191, but not Pds5-Wapl, also generated this tight DNA binding (Figure S1H), which we hereafter refer to as “DNA gripping”.

While the gripping state tolerated physiological salt washes, it was sensitive towards higher salt concentrations (500 mM NaCl), suggesting that it arises from electrostatic interactions. Such a high-salt wash removed gripped cohesin from linear DNA, but not from DNA loops with both ends tethered to the beads (Figure 1E). This suggests that, in addition to high-salt sensitive gripping, cohesin retains high-salt resistant topological association with DNA, corresponding to the previously observed topological cohesin binding using ATP ground state mimics (Çamdere et al., 2018; Minamino et al., 2018).

Lastly, we needed to know whether the gripping state is equivalent to fully topologically loaded cohesin following ATP hydrolysis. In both cases, cohesin is retained on topologically closed DNA following high-salt washes that are usually performed on ice. However, when we incubated cohesin-DNA complexes in a high salt buffer at 32 °C for 60 minutes, only cohesin that was loaded by ATP hydrolysis was retained on DNA. Cohesin in the gripping state was instead lost following this incubation (Figure S1I). Together, these data suggest that the cohesin gripping state includes topological DNA embrace, but that it is biochemically distinct from and less stable than fully topologically loaded cohesin.

### Cryo-EM Structure of Cohesin in the Gripping State

To understand the molecular architecture of the gripping state, we visualized this cohesin-DNA-loader complex by electron microscopy. We assembled cohesin onto a linear 125 bp dsDNA substrate in the presence of Mis4-Ssl3 and ADP·BeF_3_^-^ and separated the gripping reaction by sucrose gradient centrifugation (Figure 2A). The cohesin tetramer, loader and DNA co-fractionated and the peak fractions were applied to EM grids stained with uranyl acetate. Particles were homogeneously distributed and 2D averages revealed a Y-shaped complex (Figures S2A and S2B). A 3D reconstruction revealed two extended protrusions, in the shape of a bean and a rod respectively, which asymmetrically depart from the core of the complex (Figures 2B and S2C-E). A U-shaped density was apparent as part of the core, reminiscent of the *A. gossypii* and *C. thermophilum* cohesin loader subunit Scc2 (Chao et al., 2017; Kikuchi et al., 2016). Other cohesin components and the DNA were harder to assign.

**Figure 2.**
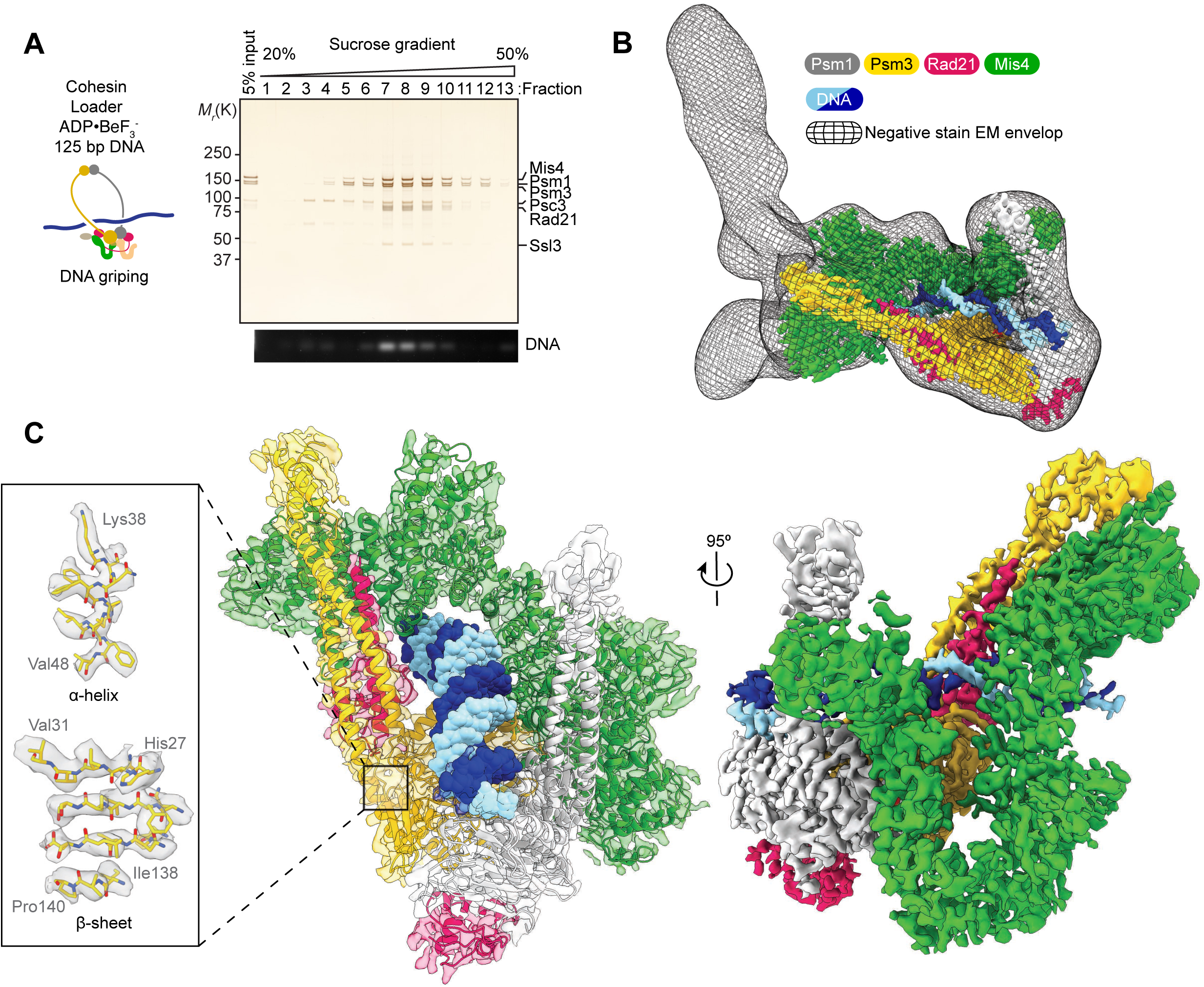
Overview Structure of Cohesin during its Loading onto DNA (A) Schematic of the EM sample preparation. A DNA gripping reaction using a 125 bp linear dsDNA substrate was separated by 25 - 50% sucrose gradient centrifugation. Protein and DNA composition of each fraction was analyzed by SDS-PAGE followed by silver staining and by agarose gel electrophoresis. Fractions 7 and 8 contained the cohesin-loader-DNA complex and were used for EM analysis. (B) Superposed image of the negative staining 3D reconstruction and the cryo-EM map of the core complex. (C) Two views of the 3.9 Å resolution cryo-EM map of the core complex with a transparent surface containing the atomic model (center) and a solid surface rendering (right). The inset on the left shows of two examples of secondary structure elements with resolved amino acidic side chains.

To visualize the cohesin-DNA complex at higher resolution, we recorded cryo-EM images of the same preparation. 2D averages revealed fine details in the core of the complex, while the protruding densities observed in the negative stain averages were less defined. Following 3D classification and local refinement, we obtained a first 3.9 Å resolution map of this core (Figures 2C and S2F – S2J). At this resolution secondary structure elements and large amino acid side chains became discernible, allowing us to build an atomic model, starting from docked homology models derived from available crystal structures of orthologous subunits (Gligoris et al., 2014; Haering et al., 2002; Kikuchi et al., 2016). The model covers the Psm1 head and proximal coiled coil bound to the Rad21 C-terminus, the Psm3 head and neck bound by the Rad21 N-terminal domain, as well as Mis4 and 32 bp of DNA.

The DNA lies on top of the engaged ATPase heads, which are nucleotide-bound and in a configuration competent for ATP hydrolysis. Mis4 clamps the DNA onto the ATPase heads, making widespread contacts with both Psm1 and Psm3. This cryo-EM volume could be docked into the negative stain map, indicating that the core of the complex overall maintains the same configuration in the room temperature-fixed sample and in the frozen-hydrated state (Figure 2B). From overlaying the two reconstructions, we see that the bean and rod in the negative stain map contact N-terminal Mis4 and the Psm3 coiled coil region.

### The Molecular Action of the Cohesin Loader

Close inspection of our new atomic model reveals two contact regions between Mis4 and cohesin. One is the U-shaped Mis4 hook that binds the engaged Psm1-Psm3 ATPase heads, and a second is the Mis4 N-terminal handle that touches the Psm3 neck region, close to the Rad21 triple helix (Figure 2C). Mis4 and Psm3 together form a protein ring that topologically encircles the DNA. The lumen of the ring is lined with positive charges, mainly clustered on the Mis4 surface (Figure 3A). A protein ring partly formed by Mis4 in the gripping state is unexpected and distinct from the well-established cohesin ring. Our new structure explains why ATP binding and not hydrolysis is sufficient for topological DNA binding (Çamdere et al., 2018; Minamino et al., 2018). Two distinct topological interactions are formed between protein and DNA during cohesin loading. One interaction is a loading intermediate, which directly involves the Mis4 loader and depends on ATP head engagement, shown here. The second interaction is the end result of the cohesin-loading reaction (Haering et al., 2008), which involves the topological entrapment in the main SMC ring and requires ATP hydrolysis.

**Figure 3.**
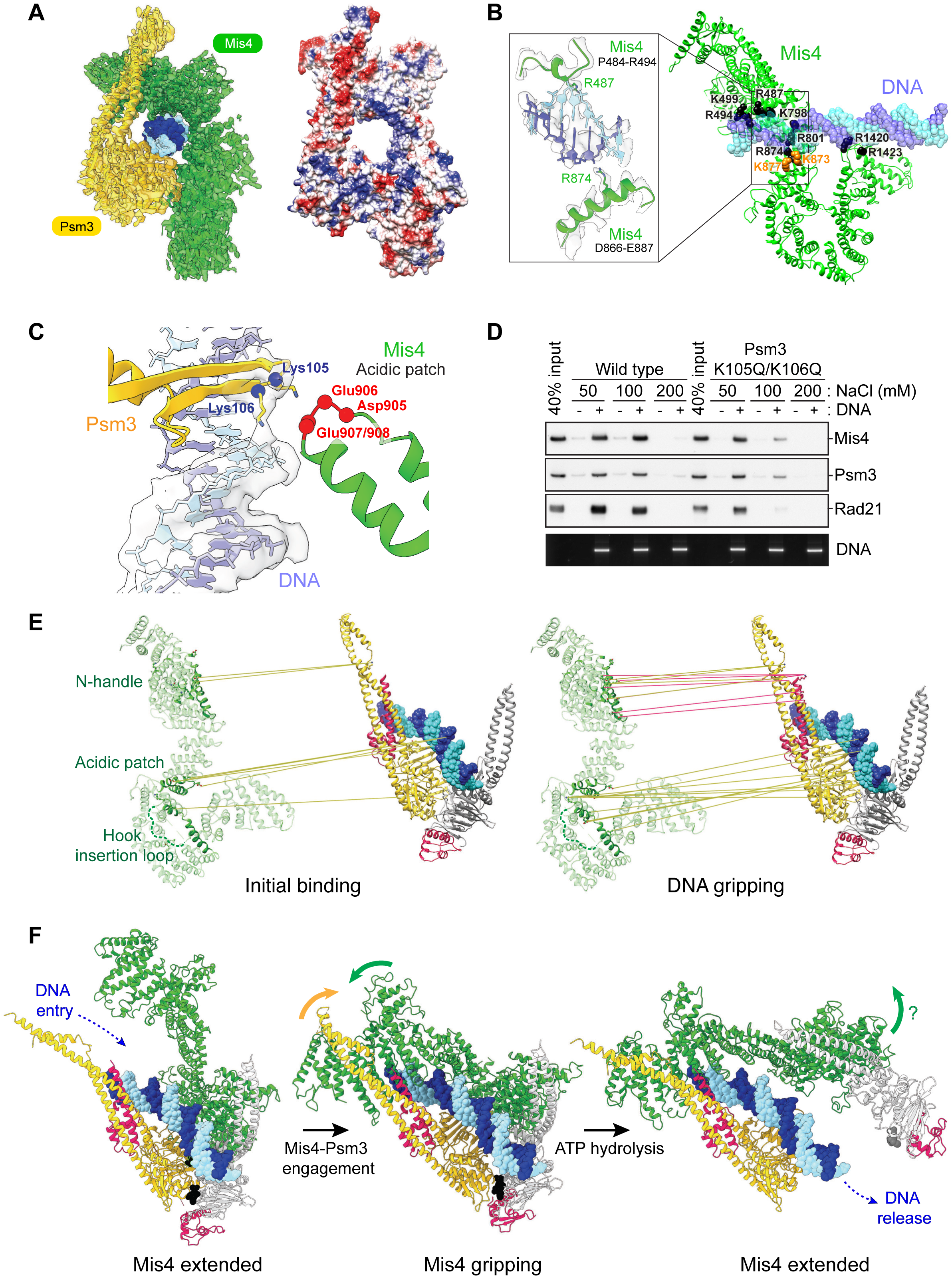
Molecular Mechanism of the Cohesin Loader (A) Psm3 and Mis4 topologically embrace DNA in the gripping state. Atomic model of Psm3, Mis4 and DNA built into the cryo-EM map (left), as well as Coulombic surface coloring for the protein component are shown. Blue represents positively and red negatively charged amino acids, respectively. (B) Positively charged residues on the Mis4 surface, colored black, line the DNA path. The inset displays the cryo-EM map and atomic model to illustrate direct contacts made by R487 and R874 with the DNA. K873 and K877, highlighted in gold, correspond to conserved *S. cerevisiae* K788 and R792, whose mutation compromises cohesin loading (Chao et al., 2017). (C) Atomic model and cryo-EM map surrounding Psm3 K105 and K106 acetyl acceptor lysines, illustrating their orientation with respect to DNA and a Mis4 acidic patch. (D) DNA gripping experiment comparing wild type and acetyl acceptor lysine mutant (K105Q/K106Q) cohesin. Following the binding incubation, the DNA beads were washed with buffer containing the indicated NaCl concentrations. (E) Comparison of CLMS contacts between initial binding and the DNA gripping state. Crosslinks between Mis4 and Psm3 (golden lines) and between Mis4 and Rad21 (red lines) were mapped onto an expanded atomic model of the DNA gripping state. Three interacting Mis4 regions are highlighted. (F) Hypothetical sequence of ATP hydrolysis-controlled Mis4 conformational changes before and after gripping state formation.

In addition to topologically entrapping DNA, Mis4 is involved in multiple DNA contacts. Regions where the Mis4 cryo-EM density is connected with DNA include R487 and R874, where the N-terminal handle and the C-terminal hook clamp the double helix (Figure 3B). Two conserved positively charged residues, K873 and K877, map in close proximity and are required for cohesin loading onto chromosomes in *S. cerevisiae* (Chao et al., 2017). The engaged SMC heads provide an additional, composite DNA binding surface, lined with conserved positively charged residues on both Psm1 and Psm3 (Figure S3A). These extensive DNA contacts likely underlie the tight electrostatic binding of linear DNA in the gripping state.

When we superimpose the engaged, DNA-bound SMC heads in our structure with the equivalent domains of Rad50 from the Rad50-Mre11-DNA complex, we find a striking overlap between both the engaged ATPase heads as well as the DNA (Figure S3A; Liu et al., 2016; Schüler and Sjögren, 2016; Seifert et al., 2016).

Furthermore, positively charged residues that make electrostatic interactions with DNA are spatially conserved on the ATPase surface, despite the limited sequence conservation between Rad50 and Psm1/Psm3. This suggests fundamental similarities in the DNA binding mechanisms of these distant relatives of the SMC family.

Amongst residues that make direct contact with the DNA is Psm3 K106, one of the conserved acetyl-acceptor lysines that convey DNA-stimulated ATP hydrolysis (Murayama and Uhlmann, 2015). Its DNA engagement is supported by connecting density in the cryo-EM map (Figure 3C). The second acetyl-acceptor, K105 does not make obvious DNA contact. Rather, cryo-EM density inspection indicates that K105 is oriented towards a conserved acidic surface loop on Mis4 (Figures 3C and S3B). Thus K105 and K106 emerge as a signaling node where DNA and the cohesin loader converge. This arrangement explains why ATPase stimulation of cohesin depends on the presence of both DNA and the cohesin loader (Murayama 2014). To investigate the contribution of the two lysines to gripping state formation, we mutated both residues to asparagine. While DNA gripping was still observed with this KKQQ mutant at low salt concentration (50 mM NaCl), contact was lost at an intermediate salt concentration (100 mM) while wild type cohesin retained tight DNA binding (Figure 3D). This observation confirms an important contribution of K105 and K106 to gripping state formation.

To further explore the Mis4-cohesin interactions, we performed protein-protein crosslinking mass spectrometry (CLMS) using a bifunctional, UV activated (NHS-diazirine) crosslinker (Figure S3C). We compared the loader-cohesin contacts in the absence and presence of nucleotide, recapitulating an ‘initial state’ before head engagement and the ‘gripping state’ described in our structure. Most crosslinks within subunits of the cohesin core (97.5%) map within 25 Å, when projected onto our cryo-EM atomic model, thus validating our approach (Figure S3D). Looking at intermolecular crosslinks, the Mis4 hook displayed numerous contacts with the Psm3 head in both states, although the identity of the amino acids involved changed between the two conditions (Figures 3E). When we map the crosslinks in the gripping state to the atomic model, we detect contacts between the Psm3 head and both flanks of the Mis4 hook. These include crosslinks with a characteristic, conserved loop that emerges from the C-terminal Mis4 flank and crosses the hook crevice to the N-terminal flank before looping backwards (Figures 3E and S3B, E). Close proximity between Psm3 K105 with the Mis4 acidic patch was also confirmed in this crosslinking experiment.

In contrast to these prevalent hook interactions, CLMS contacts of the Mis4 handle with the Psm3 neck were scarce in the initial state but became more prominent in the gripping state (Figure 3E). Conserved Mis4 handle residues (Figure S3B) make crosslinks with the Rad21 N-terminal domain in the gripping state, but these were absent in the initial state. We can rule out that lack of interactions between the Mis4 handle and N-terminal Rad21 reflect an absence of Rad21, as Rad21-Psm3 neck crosslinks were detected in both states. These observations open the possibility that Mis4 hook engagement with the Psm3 head is created in the initial state, while Mis4 interactions with the Psm3 neck and Rad21 are stabilized upon ATP-dependent DNA binding.

In our cryo-EM structure of the gripping state, Mis4 shows a striking conformational change when compared to the crystal structure of free *C. thermophilum* Scc2, the functional ortholog of Mis4 (Kikuchi et al., 2016). While the U-shaped Mis4 hook can be superimposed with a RMSD of only 1.0 Å, the angle at which the N-terminal handle emerges is tilted by 45° (Figure S3F). If we hypothetically superpose Mis4 via the U-shaped hook domain in the X-ray form to our gripping-state structure, the N-terminal handle becomes disengaged from the Psm3 neck and N-terminal Rad21, hence opening up a corridor for DNA to access the ATPase (Figure 3F, left). Given that the Mis4 N-terminal handle engages the double helix in our structure, we speculate that DNA entry itself contributes to rearranging the handle on route to gripping-state formation (Figure 3F, middle). To facilitate Mis4 engagement, the Psm3 coiled coil rotates by 25° with respect to its ATPase head, compared with other available structures of cohesin SMCs that were captured with engaged ATPase heads (Gligoris et al., 2014; Muir et al., 2020), (Figure S3G).

Mis4 and DNA contact with Psm3 K105 and K106 in the gripping state should trigger ATP hydrolysis and consequent SMC head disengagement. A model for head disengagement upon nucleotide release can be created by integrating cryo-EM and crystallographic information on Mis4. To this end, we first defined Psm3 with the N-terminal Mis4 handle and Psm1 with the C-terminal Mis4 hook as two separate rigid bodies. We then modeled a Mis4 reconfiguration from the gripping state to its extended X-ray form, which pushes the ATPase heads apart, opening a path for DNA passage through the SMC head gate (Figure 3F, right). The tendency for Mis4 to increase the Psm1-Psm3 head distance in the absence of nucleotide (Figure 1B) is consistent with this scenario.

### A Model of the Complete Cohesin Complex

To understand how loader-facilitated DNA passage through the SMC head gate contributes to topological cohesin loading, we need to place this reaction into the context of the complete cohesin complex. Our cryo-EM study yielded an atomic model of a core of the cohesin complex. To describe its complete architecture, we combined information from multibody refinement of peripheral elements, the negative stain reconstruction, as well as CLMS data to generate a hybrid model that describes cohesin in the gripping state (Figure 4A).

**Figure 4.**
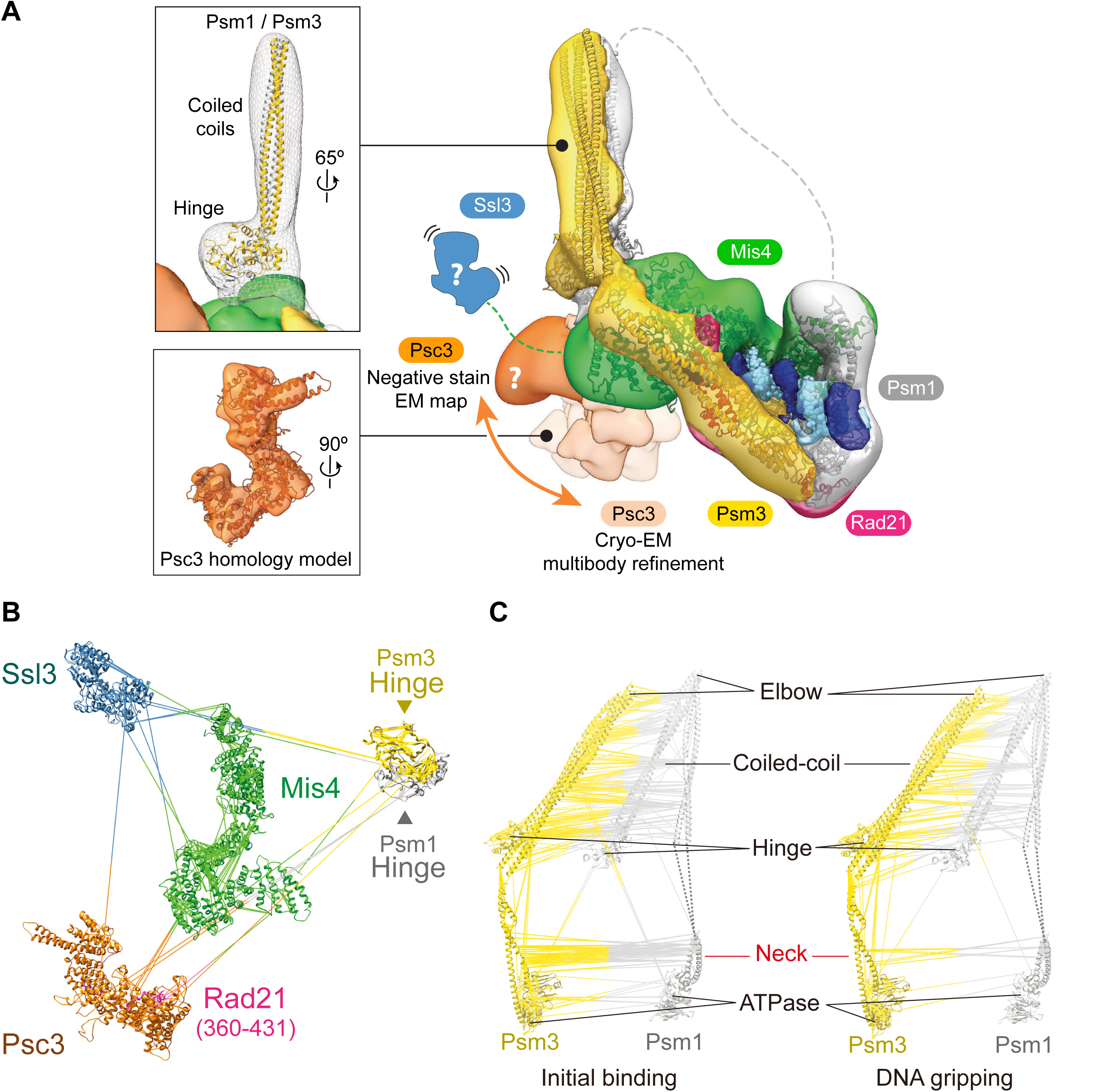
A hybrid structural model of the cohesin complex in the gripping state (A) A hybrid structural model of the cohesin complex. The atomic model of the cohesin core is docked into the negative stain EM envelope. An atomic model of the hinge and coiled coil is placed into the rod-shaped extension. The overall density accommodates a large portion of the Psm3 coiled coil, while parts of Psm1 remained invisible. The structure of Psc3 derived from multibody refinement of the cryo-EM structure is shown overlaid with the negative stain reconstruction. Variable Psc3 positions relative to the cohesin core structure are indicated, as defined by the multibody refinement analysis. Psc3 in the multibody cryo-EM structure maps in close proximity to a similarly-sized feature in the negative stain envelope. The likely position of Ssl3 bound to the Mis4 N-terminus is indicated using a cartoon representation. (B) Protein crosslinks between the atomic models in the gripping state, which support the assignments presented in panel (A). (C) Comparison of protein crosslinks within and between the SMC coiled coils in the initial binding and gripping state.

Psc3 is an essential component of the cohesin complex. By multibody refinement using masks for the core region and a peripheral, Mis4-contacting density feature, we could identify a distinct rigid body. Although only resolved to ∼10 Å, this new feature unambiguously matches a Psc3 homology model based on the *S. cerevisiae* Scc3 crystal structure (Li et al., 2018) (Figures 4A and S4A). The degree of flexibility, derived from the multibody refinement, implies a loose association with the cohesin core, at least in the state captured in our structure. In agreement with our assignment, several crosslinks in our CLMS dataset can be observed between Psc3 and Mis4 and the Psm3 head (Figure 4B and S4B). The negative stain 3D reconstruction contains a bean-shaped feature, also mapping in close proximity to the Psm3-Mis4 channel (Figure 4A). Should this feature also corresponds to Psc3, it would appear further tilted, suggesting a large degree of flexibility relative to the cohesin core.

Ssl3 is the small subunit of the cohesin loader that plays a crucial role during *in vivo* cohesin loading in the context of chromatin (Chao et al., 2015; Hinshaw et al., 2015). While Ssl3 was part of the cohesin complex in our preparation, it remained invisible in the EM structure, probably because it remains loosely tethered to the core of the complex. Nevertheless, our CLMS analysis revealed numerous crosslinks involving this subunit (Figures 4B and S4B). As expected, Ssl3 efficiently crosslinks with the Mis4 N-terminus that it encapsulates (Chao et al., 2015; Hinshaw et al., 2015). Further Ssl3 crosslinks were detected with Mis4, Psc3 and Psm3, consistent with a flexible position of Ssl3 on the posterior face of the cohesin core. Given the low resolution, we cannot exclude the possibility that the bean-shaped feature corresponds to Ssl3, not Psc3, in our negative stain reconstruction. The implications of this positioning for interactions with chromatin receptors and the *in vivo* mechanism of cohesin loading remains to be explored in the future.

A feature unique to the negative stain reconstruction is a prominent rod that projects from between the Mis4 handle and the bean-shaped feature. Its dimensions are well compatible with a model of the cohesin hinge connected to the SMC coiled coils, and atomic docking indicates that the Psm3 hinge makes direct contacts with Mis4. Crosslinks detected between the hinge and both Mis4 and Psc3 support this assignment (Figures 4A, B and S4B).

The hinge is connected with the ATPase heads via long stretches of coiled coil. Numerous intramolecular crosslinks in our CLMS dataset reflect coiled coil formation (Figures 4C and S4C). Intermolecular crosslinks between Psm1 and Psm3 suggest that both arms extend in parallel from the hinge and, in the gripping state, interact with each other up to about two thirds of their length. However, a large set of intra- and intermolecular crosslinks cannot be explained simply by an extended coiled coil configuration, as the distance between crosslinked residues largely exceeds the expected linker length. This observation indicates that the coiled coil turns back on itself with an inflection point consistent with its predicted elbow (Bürmann et al., 2019; Figures 4C and S4C, D). Consistent with coiled coil folding at the elbow, we also observed crosslinks where the hinge is expected to touch down on the coiled coil. These crosslinks all emanate from the same hinge face, suggesting that folding is directional. Based on these constraints, we can model a folded Psm3 coiled coil, showing good agreement with the negative stain envelope (Figure 4A). We did not observe continuous EM density for the Psm1 coiled coil, whose path therefore remains tentative. A single coiled coil feature is likely too thin to be visualized by negative-stain EM. In addition, SMC coiled coils are known to be flexible and to adopt a wide range of conformations (Anderson et al., 2002; Eeftens et al., 2016). Hence, our structural model reflects the observed positioning, but does not suggest a fixed orientation for the coiled coil in the gripping state.

When nucleotide was omitted in the initial state, coiled coil crosslinks were also indicative of inflection at the elbow. Contacts between the hinge and distal coiled coil further suggest that bending at the elbow is a frequent event. In addition, intermolecular Psm1-Psm3 crosslinks in the initial state extended further along the coiled coil and towards the heads (Figure 4C). This observation could indicate a defined state where the Psm1-Psm3 necks come in closer proximity. Alternatively, these Psm1-Psm3 crosslinks could report on an enhanced structural flexibility in the absence of nucleotide. In support of the latter scenario, nucleotide-dependent head engagement in the gripping state forces the Psm1 and Psm3 necks in a divergent configuration, hence precluding coiled coil interactions proximal to the heads.

### The Kleisin Path in the Gripping State

A crucial component of cohesin is its kleisin subunit Rad21 that bridges the ATPase heads. While we can see the kleisin N-terminus engaging the Psm3 neck to form a triple helix, as well as the C-terminal winged helix domain touching the Psm1 head (Figure 2C), the kleisin middle region is not resolved in our structure, as expected from the paucity of predicted secondary structure elements. To trace the kleisin path, we again turned to our CLMS analysis that contained a number of sequential crosslinks between Rad21 and the two HEAT subunits Mis4 and Psc3 (Figure 5A and S5A). As expected (Hara et al., 2014; Kikuchi et al., 2016; Li et al., 2018), Rad21 amino acids 77 – 221 line the Mis4 handle before crossing over to the C-terminal flank of the hook. The central amino acids 356 – 443 in turn follow the Psc3 body in an N- to C-terminal direction. When we project this kleisin path onto our hybrid model of the cohesin complex, this trajectory suggests that the kleisin encircles the DNA in cohesin’s gripping state (Figures 5B).

**Figure 5.**
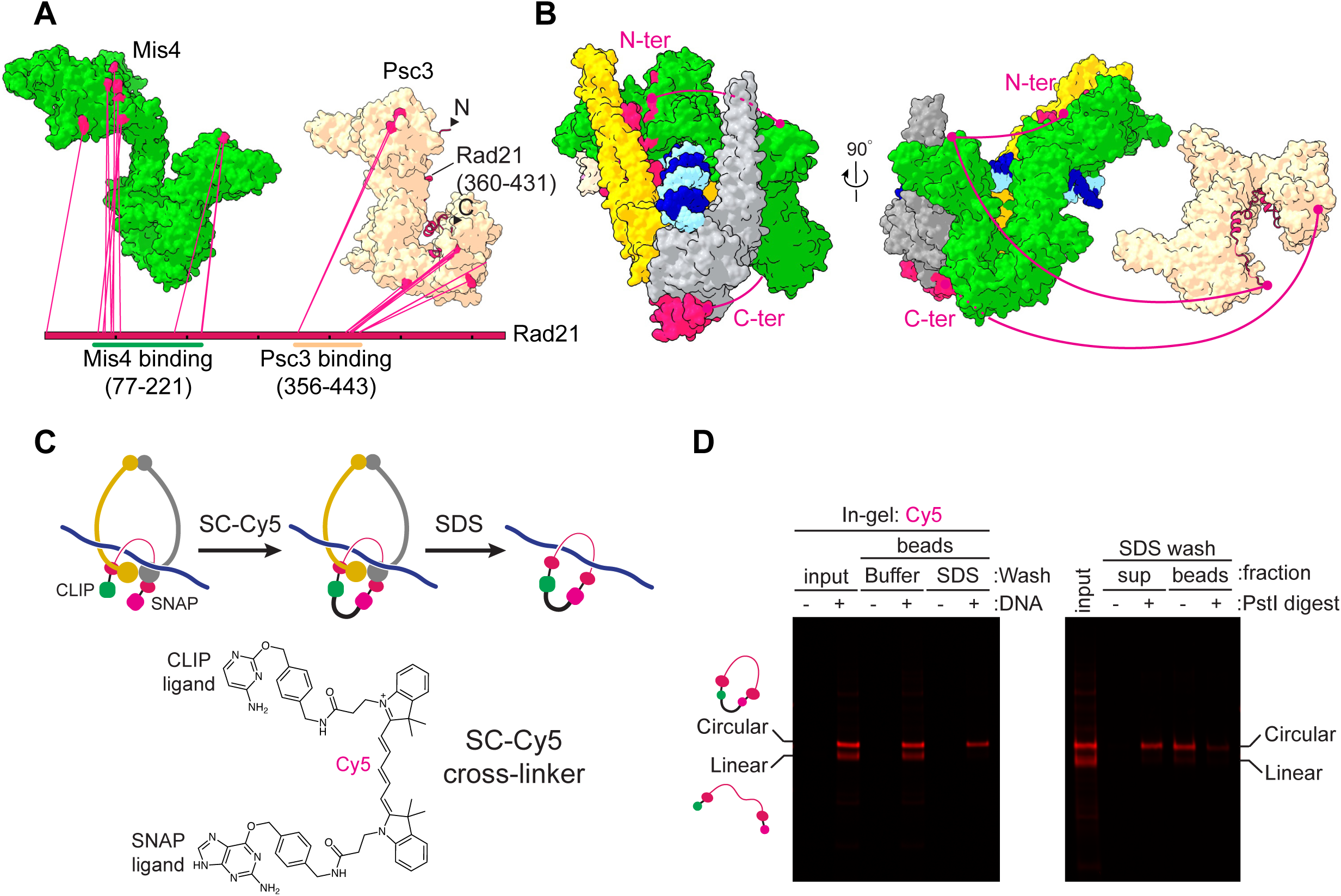
The Kleisin Path in the Gripping State (A) Crosslinks of Rad21 with Mis4 and Psc3 in the gripping state mapped onto their atomic models. Rad21 amino acids 360 – 431 are modeled based on the crystal structure of human SA2 bound to Rad21 (PDB: 4PK7) (Hara et al., 2014). (B) The crosslink sites from panel (A) are mapped on the structures of gripping state complex components, suggesting a likely kleisin path (red line). (C) Schematic of the kleisin circularization experiment. A gripping reaction was performed with Rad21 carrying N- and C-terminal CLIP and SNAP tags using a DNA loop substrate on beads. The CLIP and SNAP tags were now covalently crosslinked by SC-Cy5 (its chemical structure shown in the bottom panel). If the predicted kleisin path is correct, the kleisin protein will remain topologically associated with the DNA following denaturation with SDS. (D) In gel Cy5 detection of the experiment in (C). SC-Cy5 was added to the input proteins or following gripping state assembly on the DNA beads. Beads were then washed either with buffer or with SDS (left). After the SDS wash, DNA beads were treated with or without PstI restriction endonuclease and bead-bound and supernatant fractions analyzed (right)

To probe this suggested kleisin topology with respect to the DNA, we covalently joined the kleisin N and C termini in the gripping state. This should result in topological DNA entrapment by the kleisin (Figure 5C). We fused the kleisin to CLIP and SNAP tags at the N and C termini, respectively (Figure S5B). To covalently link the two tags, we chemically synthesized a crosslinker in which a SNAP substrate is linked to a CLIP substrate via a Cy5 dye moiety (SC-Cy5, Figure 5C). We then carried out a DNA gripping reaction using a DNA loop substrate, attached to magnetic beads. Following the gripping reaction, we added SC-Cy5. This resulted in approximately equal proportions of linear Rad21, labelled at one or both ends, and circularized Rad21. The latter was identified by its retarded gel mobility (Figure 5D and S5C). Cohesin was then denatured in buffer containing SDS, to assess whether the kleisin was retained on the DNA. Circularized Rad21, but not linear Rad21 or Psm3, remained bound to the DNA beads. The topological nature of circularized Rad21 binding to the DNA was confirmed by cleaving the DNA with the PstI restriction endonuclease, which resulted in Rad21 elution from the beads (Figure 5D). This result confirms that the kleisin indeed encircles the DNA in the gripping state. We used a similar approach with two pairs of SNAP and CLIP tags to covalently close both the SMC heads and hinge simultaneously. This experiment revealed that DNA is not yet entrapped within the circularized SMC ring (Figures S5D-F).

### The DNA Trajectory into the Cohesin Ring

While our cryo-EM structure describes the double helix entrapped between the kleisin and ATPase head gate, it does not discriminate between two possible DNA access routes. Does the DNA enter from the bottom of the ATPase, having passed through the head gate, or rather from the top of the ATPase and through the kleisin N-gate? By inspecting our structure alone, we can also not exclude that a short duplex DNA segment occupying the Mis4-Psm3 cavity is an *in vitro* artifact. To explore how DNA reaches the gripping state, we developed a protocol for DNA-protein crosslink mass spectrometry (DPC-MS, Figure 6A).

**Figure 6.**
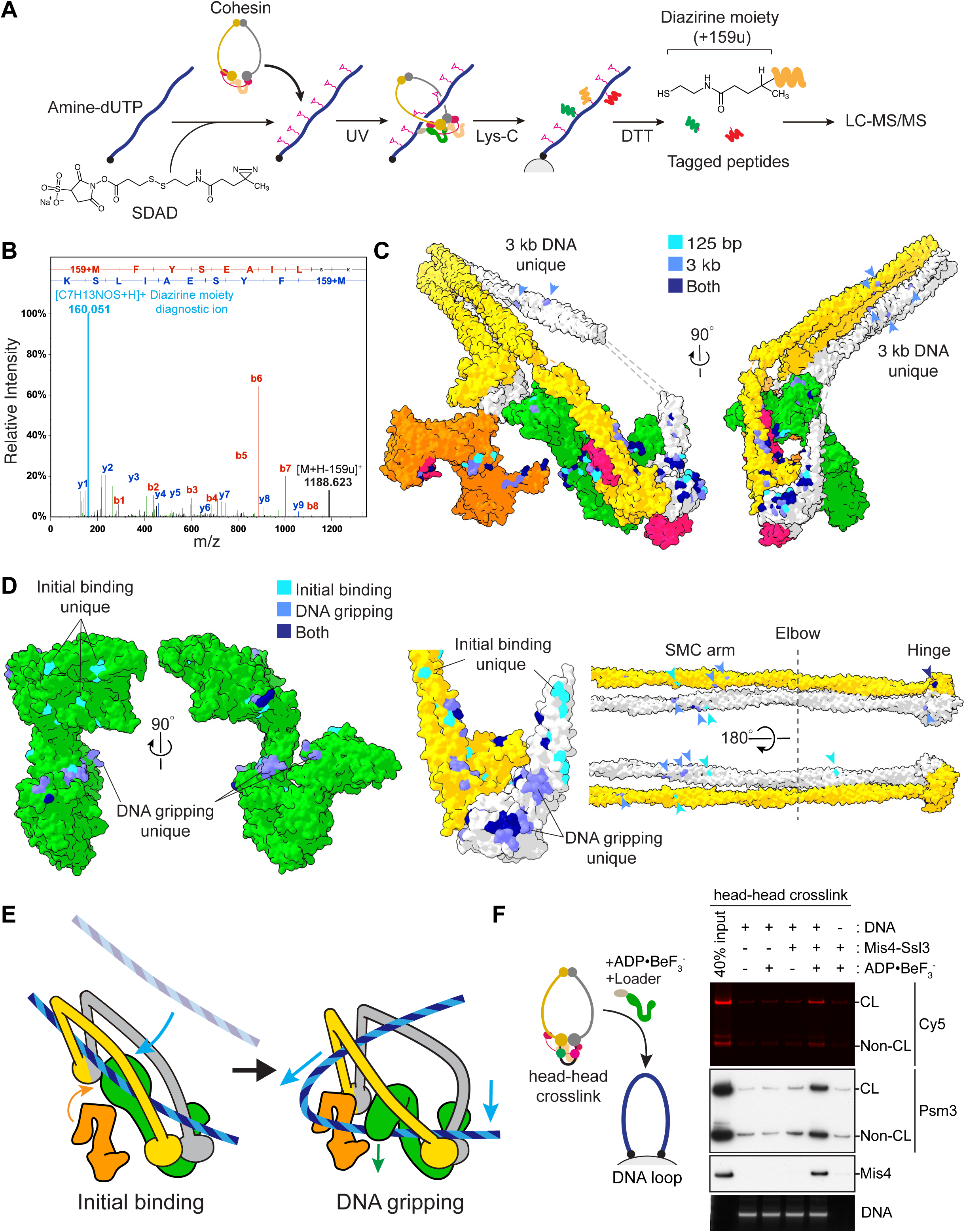
The DNA Trajectory Into the Cohesin Ring (A) Schematic of the DNA-protein crosslinking mass spectrometry workflow. See the main text for details. (B) A representative mass spectrum of a peptide containing a diazirine mass tag transferred from DNA. The diagnostic 159u ion is highlighted. (C) DNA crosslinks of a 125 bp linear DNA in the gripping state shown on the surface of the hybrid model (light blue), compared to the crosslinks observed with a 3 kb circular plasmid DNA (medium blue). Crosslinks in common to both are shown in dark blue. (D) DNA crosslinks in the initial DNA binding state (light blue) are compared to those in the gripping state (medium blue), while those in common are shown in dark blue. Surface representations of Mis4 and the SMC subunits are shown. Arrowheads highlight crosslinks along the SMC coiled coils and hinge. (E) A model of the DNA trajectory from initial binding toward the gripping state, based on the observed DNA contacts. (F) DNA gripping experiment using head-head crosslinked cohesin. Psm1-SNAP Psm3-CLIP cohesin was treated with SC-Cy5 to close the head gate before the gripping reaction using a bead-bound DNA loop substrate. DNA-bound proteins were analyzed by immunoblotting and in gel Cy5 detection.

To mark the DNA binding site on a protein, we designed a photo-crosslinkable DNA probe. First, amine-dUTP is incorporated, in place of dTTP, during DNA synthesis. The amino groups are then decorated with a bifunctional succinimidyl-SS-diazirine (SDAD) crosslinker. Nucleoprotein complexes are assembled with this probe and UV-irradiated to induce DNA-protein crosslinking. Proteins are then digested using the Lys-C endopeptidase and the DNA covalently linked to peptide fragments is isolated by biotin affinity pulldown. Finally, the SDAD crosslinker is cleaved at its disulfide bond under reducing conditions to elute the recovered peptides. These proteolytic fragments retain a characteristic mass tag of +159u at the crosslink position that can be identified by LC-tandem MS, mapping next to a diagnostic 159u peak that stems from loss of the mass tag upon peptide fragmentation (Figure 6B).

We first performed DPC-MS analysis using a derivatized 125 bp linear DNA probe, based on our cryo-EM structure. The observed DNA-protein crosslinks identified surface exposed amino acids that map in proximity to the double helix in our structure along the Mis4 hook, on the Psm3 and Psm1 heads, as well as Psc3 (Figures 6C and S6A).

Next, we repeated the DPC-MS analysis of the gripping state with a 3 kb covalently closed circular dsDNA as the substrate. Comparison with the short linear DNA revealed a near identical range of DNA-protein crosslinks (Figure 6C and S6A). We conclude that the DNA position observed in our cryo-EM structure is a fair reflection of that reached by a topologically closed DNA that reflects a more natural substrate. Additional DNA contacts were observed with long circular DNA involving the Psm1 and Psm3 coiled coil. These contacts inform us on a likely path that a longer DNA takes in the gripping state (see below).

To understand the trajectory taken by DNA to reach the gripping state, we compared the protein crosslinks with a long circular DNA probe in the initial binding state where ATP is omitted, and the nucleotide-bound gripping state. This revealed notable differences. Unique DNA crosslinks were identified in the initial state that map far above the ATPase head, while the ATPase heads are a major DNA-interaction site in the gripping state (Figures 6D and S6B). On Mis4, initial state crosslinks map on top of the handle, while in the gripping state the crosslinks line the hook. These observations suggest that DNA reaches the gripping state approaching Mis4 and the ATPase heads from the top and not the bottom.

We also recorded several DNA contacts along the SMC coiled coils and hinge in both the initial and the gripping state. All hinge crosslinks map on a solvent-exposed hinge surface opposite to that engaged in protein-protein contacts with the cohesin core (Figures 6D). These interactions further support a likely DNA entry path from the top of the ATPase that is schematically depicted in Figure 6E.

Our proposed model for DNA entry makes the prediction that DNA would not traverse the head gate on its way to the gripping state. To test this prediction, we prepared cohesin with SNAP and CLIP tags fused to the Psm3 and Psm1 C-termini, respectively. SC-Cy5 addition yielded covalent head gate closure in approximately half of the cohesin population (Figure S6C). This mixture was employed in a DNA gripping reaction using a DNA loop substrate on magnetic beads. This revealed that the efficiency of DNA engagement remained unchanged irrespective of whether the SMC head gate was open or closed (Figure 6F). This observation suggests that the DNA does not need to traverse the SMC head gate for DNA gripping to occur, further supporting DNA access from the top of the ATPase.

### ATP-dependent Kleisin N-gate Opening

If DNA accesses the ATPase from the top, then the kleisin N-gate must open to let the DNA enter, before the kleisin can encircle the DNA, as observed in the gripping state. To understand how the kleisin N-gate is regulated, we performed FRET measurements between donor and acceptor fluorophore-labeled CLIP and SNAP tags attached to the Psm3 and Rad21 N-termini (Figures 7A and S7A). A cohesin tetramer in the incubation buffer displayed measurable FRET, consistent with proximity. Addition of ATP, with or without DNA and loader, led to FRET loss, suggestive of kleisin N-gate opening. In contrast, addition of DNA or loader in the absence of ATP did not induce a FRET change. Only during gripping state formation in the presence of the Mis4-Ssl3 loader, DNA and non-hydrolyzable ATP, we observed a distinct FRET increase, consistent with kleisin N-gate closure in this state. These observations suggest that ATP binding promotes kleisin N-gate opening, but that the gate is closed again when the loader clamps DNA upon gripping state formation.

**Figure 7.**
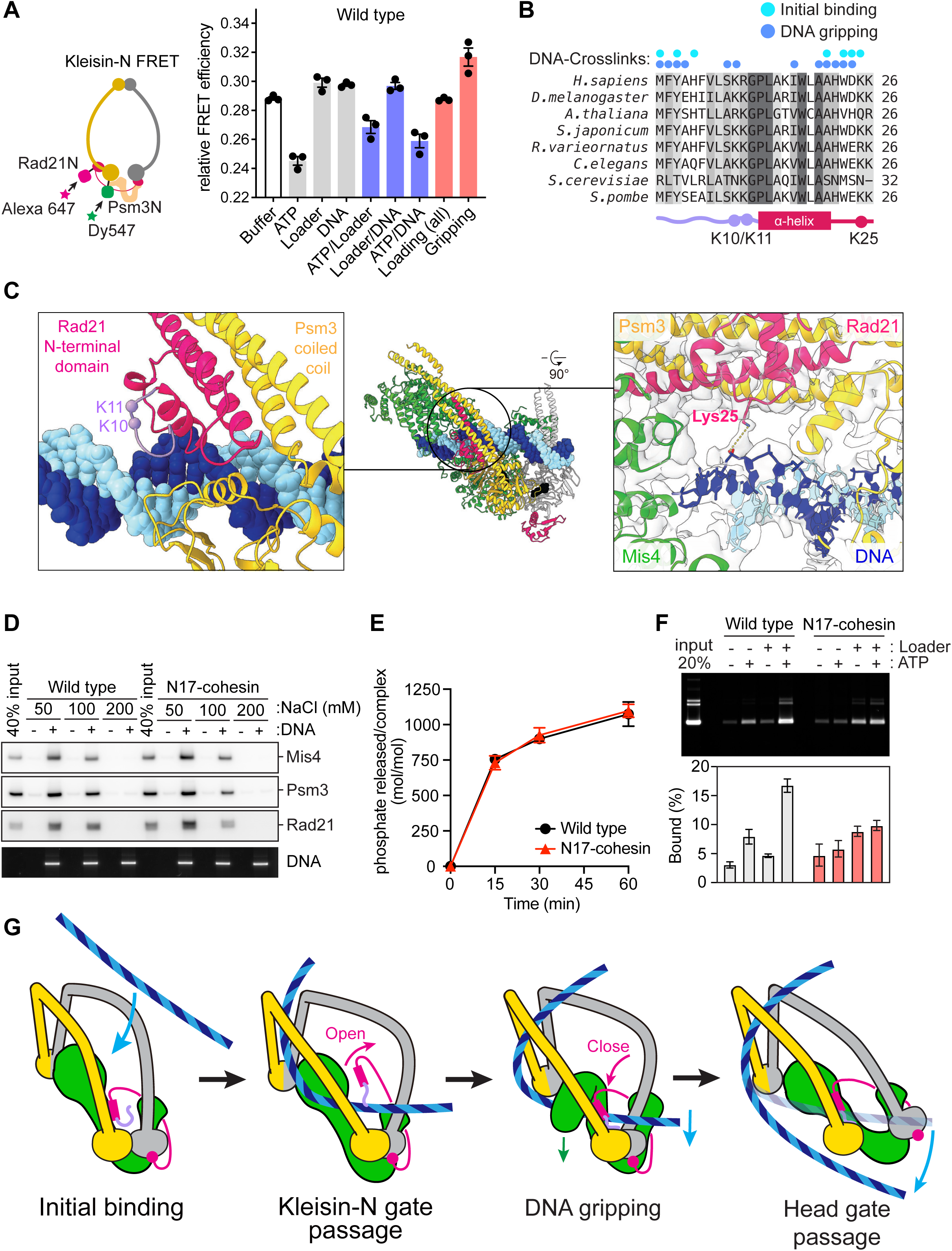
A Kleisin N-terminal Tail Guides DNA into the Cohesin Ring (A) Schematic of the kleisin N-gate FRET construct. A SNAP-tag on the kleisin N-terminus and CLIP-tag on the Psm3 N-terminus were labeled with Alexa 647 and Dy547, respectively. FRET efficiencies at the kleisin N-gate were recorded under the indicated conditions using a 3 kb plasmid DNA as a substrate. ADP▪BeF^3-^ was used as the nucleotide in the gripping incubation. Results from three independent repeats of the experiment, their means and standard deviations are shown. (B) Sequence alignment of the cohesin N-terminus. Conservation is marked by shades of grey. Positions of DNA crosslinking in the initial binding and the DNA gripping state are indicated. (C) Atomic model of the Rad21 N-tail (left), showing the position of the conserved K10 and K11 residues relative to the DNA. A magnified view around K25 is also shown (right), including the cryo-EM density that demonstrates DNA contact. (D) Comparison of wild type and N17-cohesin in a DNA gripping experiment. Following reaction with a bead-bound DNA loop substrate, the beads were washed with buffer containing the indicated NaCl concentrations. Bead-associated proteins and DNA were analyzed by immunoblotting and gel electrophoresis. (E) Comparison of ATP hydrolysis by wild type and N17-cohesin in the presence of loader and a 3 kb plasmid DNA. Shown are the means and standard deviations from three independent experiments. (F) Loading of wild type and N17-cohesin onto a 3 kb plasmid DNA in the presence of loader and ATP. Following the loading reaction, cohesin was immunoprecipitated, washed with buffer containing 750 mM NaCl and recovered DNA was analyzed by agarose gel electrophoresis. Shown are the means and standard deviations from three independent experiments. (G) A model for DNA entry into the cohesin ring. The kleisin N-tail guides DNA through the kleisin N-gate before DNA reaches the ATPase heads. ATP hydrolysis and passage through the head gate completes DNA entry.

To evaluate how ATP opens the kleisin N-gate, we asked whether ATPase head engagement or ATP hydrolysis is required in this process. To this end, we repeated the FRET experiment using signature motif mutant of cohesin, defective in head engagement (Hopfner et al., 2000). This mutant markedly dampened the FRET loss following ATP addition (Figure S7B). On the other hand, EQ-cohesin, defective in ATP hydrolysis, displayed reduced FRET and recapitulated ATP-dependent kleisin N-gate opening. Our observations suggest that ATPase head engagement triggers kleisin N-gate opening, but that the gate closes again as DNA and the cohesin loader assemble to form the gripping state. These conclusions are supported by recent observations of an open kleisin N-gate in structures of the head-engaged cohesin and condensin ATPase (Hassler et al., 2019; Muir et al., 2020).

### The Kleisin N-terminus Guides DNA into the Cohesin Ring

Evidence acquired so far indicates that DNA must diffuse through the kleisin N-gate before reaching the gripping state. However, merely based on our structure, it is unclear what would prevent DNA from reaching the gripping state without traversing the kleisin N-gate (Figure S7C). Our attention was drawn to the extreme N-terminal 12 amino acids of Rad21 that precede the alpha helix that takes part in triple helix formation at the Psm3 neck. This N-tail was crosslinked with DNA in our DPC-MS experiments, both in the initial as well as in the gripping state (Figure 7B). The N-tail is conserved amongst cohesins throughout evolution, including a series of positively charged residues. Kleisins from other SMC complexes, including prokaryotic members, also possess N-tails enriched for positive charges (Figure S7D). The cryo-EM map was of sufficient quality to build amino acids 5-12 of the kleisin N-tail, forming a loop wedged between the Psm3 ATPase head and the DNA. Two conserved lysines 10 and 11 point toward the DNA. While their distance is too far to maintain direct DNA contact in the gripping state, lysine 10 is amongst the residues crosslinked with the DNA (Figure 7C, left). Another conserved positive residue following the triple helix, lysine 25, directly engages DNA (Figure 7C, right). The N-tail is held in place by an extended loop projecting from the Psm3 ATPase head that is specific to this subunit and much shorter in Psm1 (Figure S7E). This observation provides further indication that SMC proteins evolved to asymmetrically interact with the kleisin subunit in cohesin.

If the kleisin N-tail binds DNA when the kleisin N-gate opens, and maintains DNA contact when the Rad21-Psm3 triple helix structure reforms, this will have guided the DNA through the kleisin gate on its way to the gripping state (Figure S7C). To analyze the contribution of the kleisin N-tail to cohesin function, we purified a cohesin complex lacking Rad21 amino acids 1-17 (N17-cohesin; Figure S7F).

When we included N17-cohesin in a DNA gripping experiment, this complex bound DNA similarly tightly compared to wild type cohesin (Figure 7D). Likewise, the levels of DNA and loader-stimulated ATP hydrolysis were equal when using wild type or N17-cohesin (Figure 7E). This suggests that the kleisin N-tail affects neither the tight DNA binding associated with gripping-state formation, nor cohesin loader and DNA-stimulation of ATP hydrolysis.

We then investigated the contribution of the kleisin N-tail to topological cohesin loading onto DNA. Following incubation in the presence of loader and ATP, we recovered cohesin by immunoprecipitation and assessed topological, high-salt resistant, DNA binding. N17-cohesin showed a substantially reduced ability to retain DNA, compared to wild type cohesin, indicating that topological cohesin loading onto DNA was unsuccessful (Figure 7F). From our results we conclude that the Rad21 N-tail has a crucial role in guiding DNA to successful DNA entry into the cohesin ring (Figure 7G).

## DISCUSSION

We have used a combination of biochemical and structural approaches to learn how the DNA enters into the cohesin ring. FRET measurements at the cohesin ATPase heads led us to discover a DNA gripping intermediate that forms when cohesin comes together with the loader, DNA and non-hydrolyzable ATP. The gripping complex proved to be a suitable target for cryo-EM imaging and allowed us to generate an atomic model of key elements of this cohesin loading intermediate. To understand where this intermediate lies during the cohesin loading reaction onto DNA, we utilized additional biochemical tools, including a newly developed DNA-protein crosslink mass spectrometry assay. These approaches allowed us to trace the DNA trajectory into the cohesin ring. We also identified previously uncharacterized functional elements of cohesin that are important for successful topological entry. Amongst these, the kleisin N-tail opens up a dichotomy of the DNA entry reaction that might be important for understanding cohesin’s alternative role in loop extrusion.

### The DNA Trajectory into the Cohesin Ring

In the gripping state, we find the DNA trapped between two gates, the kleisin N-gate and the ATPase head gate, that lead into the cohesin ring. Given this topology, the DNA must have passed one of the two gates, but not yet the other. Numerous lines of evidence indicate that the DNA arrived from the top of the ATPase and that it must have passed the kleisin N-gate. The evidence includes direct observation of kleisin N-gate opening, as well as protein-protein and DNA-protein proximity mapping before and upon gripping state formation. If DNA has passed the kleisin N-gate on its way to the gripping state, subsequent passage through the head gate must be required to complete topological entry. While we have not yet directly observed this step, the ATPase heads in the gripping state are ATP-bound and appear competent for ATP hydrolysis, which is expected to trigger head gate opening (Hopfner et al., 2000).

Using a FRET-based assay, we established that kleisin N-gate opening required ATP-dependent SMC head engagement, consistent with recent structural observations of engaged cohesin and condensin ATPase heads with an open N-gate (Hassler et al., 2019; Muir et al., 2020). How ATP binding contributes to opening of this gate remains to be fully understood. Our cryo-EM structure in turn indicates that, by the time DNA reaches the gripping state, the kleisin N-gate is closed again. In this configuration, the DNA itself contributes to keeping the gate shut by directly contacting the kleisin N-terminal domain and locking it into the triple helix with the Psm3^Smc3^ neck. ATP hydrolysis then leads to ATPase head gate opening, while the kleisin N-gate remains shut. This structural model provides an explanation for two interlocking gates through which DNA enters the cohesin ring, only one of which can be open at any one time. We previously hypothesized that DNA enters the cohesin ring through the interlocking kleisin N- and head gates (Murayama and Uhlmann, 2015). Our new molecular knowledge now allows us to establish the correct order of events that lead to DNA loading.

An alternative model for DNA entry into the cohesin ring states that DNA passes a ‘hinge gate’ (Buheitel and Stemmann, 2013; Gruber et al., 2006). This model is based on two observations. Firstly, DNA entry into the cohesin ring was blocked by ligand-induced dimerization of ectopic domains, inserted into both halves of the hinge. This result was interpreted to demonstrate DNA passage through a hinge gate. However, our structure opens up an alternative explanation. We find that the hinge makes contact with both the Mis4^Scc2/NIPBL^ and Psc3^Scc3/STAG1/2^ subunits in the gripping state, as well as the coiled coil arms. These contacts promote coiled coil folding at the elbow, which is crucial for DNA access to the kleisin N-gate. The hinge insertions might have interfered with one or more of these interactions, thereby compromising DNA access to the kleisin N-gate. A second result was at first sight incompatible with DNA entry through the kleisin N-gate. An Smc3-kleisin fusion protein, in which the two subunits cannot separate, remained able to load onto chromosomes (Buheitel and Stemmann, 2013; Gruber et al., 2006). However, the linkers used to connect Smc3 and the kleisin, used in these studies, are relatively long. The Smc3-kleisin fusion in itself will not block the operation of the kleisin N-gate or subsequent DNA passage through the head gate. Indeed, a functional kleisin N-gate remains required for the viability of the fusion strains (Guacci et al., 2019).

We therefore suggest that Smc3-kleisin fusion does not impede the loading reaction, it rather results in a loading product in which the linker sequence adds an additional DNA embrace. This possibility might explain the compromised function of the fusion protein (Gruber et al., 2006). Future experiments are required to clarify the topology of the DNA loading products obtained with an Smc3-kleisin fusion protein.

### The Role of the Cohesin Loader and of other HEAT Repeat Subunits

The Mis4^Scc2/NIPBL^ cohesin loader was thought to associate with the cohesin complex through kleisin interactions (Kikuchi et al., 2016). We now find that the loader engages in additional extensive interactions with both SMC subunits and the DNA. These contacts aid the topological loading reaction in several ways. When compared to its previously reported extended conformation, the loader in the gripping state has undergone a striking conformational change, contributing to the topological enclosure that holds DNA locked against the ATPase heads. Together with the DNA, the loader also engages the Psm3^Smc3^ acetyl acceptor lysines, to trigger ATP hydrolysis and SMC head gate opening. Following ATP hydrolysis, we propose that the loader might return to its extended form, further promoting head separation and DNA passage through the head gate (Figure 3F). Despite these multiple ways in which the loader facilitates DNA entry, cohesin retains basal topological loading potential without the loader (Murayama and Uhlmann, 2014). Indeed, our FRET results indicate that Kleisin N-gate opening by head engagement is independent of the loader. In this scenario, arrival of the DNA alone might be sufficient to close the N-gate, before ATP hydrolysis completes DNA entry at a reduced rate.

In addition to the Mis4^Scc2/NIPBL^ loader, the Psc3^Scc3/STAG1/2^ subunit is instrumental for cohesin loading onto DNA, both *in vivo* and *in vitro* (Murayama and Uhlmann, 2014; Tóth et al., 1999). Our structure shows that Psc3^Scc3/STAG1/2^ is positioned behind the loader in the gripping state. Furthermore, we observe DNA crosslinks, consistent with interactions that were observed crystallographically (Li et al., 2018), both in the initial DNA binding as well as the gripping state. Based on this positioning, we propose that Psc3^Scc3/STAG1/2^ plays a role in attracting DNA, as it approaches between the coiled coil arms and towards the ATPase (Figure 6E). We note that, apart from contacts with the kleisin (Haering et al., 2002; Hara et al., 2014; Li et al., 2018), the only other biochemically documented interaction of Psc3 with the cohesin complex lies at the hinge (Murayama and Uhlmann, 2015). The molecular mechanism by which Psc3^Scc3/STAG1/2^ contributes to cohesin function remains to be further explored.

Cohesin’s third HEAT repeat subunit, Pds5, is thought to replace Mis4^Scc2/NIPBL^ following loading (Murayama and Uhlmann, 2015; Petela et al., 2018). Pds5 has a similar overall shape to that of Mis4 (Lee et al., 2016; Ouyang et al., 2016). The conserved Psm3 contacting residues in Mis4’s handle are also found in Pds5 (Figure S3B), suggesting that aspects of its engagement with cohesin are likely conserved. However, unlike the loader, Pds5 does not stimulate ATP hydrolysis by cohesin (Murayama and Uhlmann, 2015) and our FRET results indicate that Pds5 fails to promote ATPase head engagement in the presence of DNA and non-hydrolyzable ATP. Instead, we speculate that Pds5 might block head engagement, thereby preventing spontaneous kleisin N-gate opening. Assuming that the DNA exit path could follow similar sequential passage through the kleisin N- and head gates, this model could explain the role of Pds5 in stabilizing cohesin on DNA. Should kleisin N-gate opening through head engagement be no longer possible in the presence of Pds5, an alternative way to operate the kleisin N-gate would become necessary.

This could be the role of Wapl that is directly recruited by Pds5. This scenario provides a rationale for how Pds5 could both stabilize cohesin on chromosomes and render it competent for Wapl-regulated unloading. How Pds5 differs from the Mis4^Scc2/NIPBL^ loader, and how their alternating association with the cohesin complex is controlled, remains to be investigated.

### The Kleisin N-tail and Its Implication for Successful DNA Entry

While performing DNA-protein crosslink experiments, we noticed crosslinks with a conserved kleisin N-tail that has previously received little attention. This tail lies in close proximity of the Psm3 head when the N-gate is closed upon triple helix formation with the Psm3 neck. If the N-tail were to maintain DNA contact during the transition from an open to a shut kleisin N-gate, the tail would have guided the DNA through the kleisin gate. In support of this notion, while dispensable for gripping-state formation and ATP hydrolysis, we find that the N-tail is key to successful DNA entry into the cohesin ring.

Why is a kleisin N-tail required to guide DNA through the kleisin N-gate? The structured components of the DNA gripping state have no obvious mechanism for sensing whether DNA has passed the kleisin N-gate or not (Figure S7C). Only our kleisin circularization experiment revealed that the DNA has in fact traversed the kleisin N-gate. We therefore speculate that, under certain conditions, DNA might reach the gripping state without having passed through the kleisin N-gate. One example is a cohesin complex lacking the N-tail. In this case, acetyl-acceptor lysine engagement by DNA and the loader still triggers ATP hydrolysis, but the outcome of head disengagement will be diametrically different: without having passed the kleisin N-gate first, DNA cannot enter the cohesin ring. Following ATP hydrolysis, the loader might dissociate or revert to its extended configuration, both of which would alter interactions of the hinge with the cohesin core and favor a transition from bent to straight SMC coiled coils. Provided Psc3 remains associated with the hinge and DNA during this process, cohesin would nucleate a DNA-loop (Figure S7C). Were such abortive DNA entry reactions to repeat, this would lead to expansion and extrusion of the loop.

In prokaryotic and eukaryotic SMC complexes, the kleisin N-tail is broadly conserved and universally enriched for positively charged amino acids. This conservation suggests that a similar dichotomy may exist in other SMC complexes, in which the kleisin N-tail mediates kleisin N-gate passage and dictates whether topological DNA entry is achieved or a loop is extruded. Whether DNA can reach the gripping state without passing the N-gate under physiological conditions remains an important question to explore.

## Acknowledgments

We would like to thank Simone Kunzelmann, Andrew Purkiss and Phil Walker for their help, Lutz Fischer for custom software, Christos Savva for support with Krios K3 image acquisition, the Crick Fermentation Science Technology Platform, as well as the members of our laboratories for valuable discussions and comments on our manuscript. This project received funding from the European Research Council (ERC) under the European Union’s Horizon 2020 research and innovation program (grant agreement Nos. 670412 and 820102) and the Francis Crick Institute, which receives its core funding from Cancer Research UK (FC001198 and FC001065), the UK Medical Research Council (FC001198 and FC001065), and the Wellcome Trust (FC001198 and FC001065). JR acknowledges funding by the Deutsche Forschungsgemeinschaft (DFG) under Germanýs Excellence Strategy-EXC 2008-390540038-UniSysCat.

## Author Contributions

T.L.H. initiated the study and performed all biochemical experiments in consultation with F.U.. P.E. performed all negative stain EM, cryo-EM image processing advised by A.C. and prepared the structural figures with T.H. and A.C.. J.L. prepared and screened cryo-EM grids. A.N. set up the Krios Falcon 3 data collection with the help of P.E. and helped with cryo-EM image processing. J.S.S. built the atomic model with the help of P.E.. H.R.F. and A.P.S. analyzed the DPC-MS samples. G.P. and N.O‘R. synthesized SC-Cy5. Z.A.C., F.J.O’R. and J.R. conducted the CLMS analysis. T.L.H., A.C. and F.U. wrote the manuscript with input from all coauthors.

## MATERIAL AND METHODS

### Yeast Strains

All fission yeast cohesin tetramer complexes and Pds5 were expressed in W303 background budding yeast strains. Strains were cultured at 30 °C in YP medium (2% peptone and 1% yeast extract) containing 2% raffinose until the optical density at 600 nm reached 1.0. Protein expression was induced by addition of 2% galactose for 4 hours. Fission yeast Mis4-Ssl3 protein or Mis4-N191 protein was expressed in fission yeast strains. Fission yeast cells were cultured in EMM minimal medium supplemented with 30 μM thiamine at 30 °C until the optical density at 595 nm reached 1.5, and protein expression was induced in EMM minimal medium lacking thiamine for 15 hours. Genotypes of all strains used are listed in Table S1.

### Bacteria

Fission yeast Wapl was expressed in the *E. coli* strain BL21-CodonPlus (DE3)-RIPL (Agilent Technologies). The genotype is: *E. coli B F-ompT hsdS(rB-mB-) dcm+ Tetr gal λ(DE3) endA Hte [argU proL Camr] [argU ileY leuW Strep/Specr]*.

### Cloning of cohesin and its variants for protein purification

For construction of Head FRET wild type and EQ-cohesin, SNAP-tag and CLIP-tag encoding sequences were fused to Psm1 C-terminus and Psm3 C-terminus in the shuttle vector YIplac211-Psm1/Psm3 or YIplac211-Psm1 _E1161Q_ /Psm3 _E1128Q_ that were constructed previously (Murayama and Uhlmann, 2014, 2015). The YIplac211-Psm1-SNAP/Psm3-CLIP vector and a YIplac128-Rad21/Psc3 expression vector were sequentially integrated into budding yeast at the *URA3* and *LEU2* loci, respectively.

For construction of the Kleisin-N FRET cohesin complex, SNAP-tag and CLIP-tag sequences were fused to Rad21 N-terminus in the YIplac128-Rad21-Psc3 integration vector and Psm3 N-terminus in the YIplac211-Psm1-Psm3 vector. Both vectors were integrated into budding yeast genome as before. Kleisin-N FRET Walker B motif mutant (EQ) and signature motif mutant (SQ) complexes were generated by site-directed mutagenesis on the YIplac211-Psm1/CLIP-Psm3 vector.

For construction of the kleisin circle construct, SNAP and CLIP-tag sequences were fused to Rad21 C-terminus and N-terminus in the YIplac128-Rad21-Psc3 integration vector. The YIplac128-CLIP-Rad21-SNAP/Psc3 expression vector was integrated into budding yeast harboring YIplac211-Psm1/Psm3.

For construction of SMC circle construct, SNAP-tag sequences were integrated into the Psm1 hinge region (between R593 and G594) and the Psm3 C-terminus. CLIP-tag sequences were integrated into the Psm3 hinge region (between S631 and N632) and fused to the Psm1 C-terminus in the YIplac211-Psm1/Psm3 vector. The expression vector was integrated into budding yeast harboring YIplac128-Rad21/Psc3.

For construction of N-terminally truncated N17-Rad21, a partial coding sequence (amino acids 18-646) was amplified by PCR, which replaced the full length Rad21 gene in the YIplac128-Rad21/Psc3 vector by In-Fusion cloning. The YIplac128-N17-Rad21/Psc3 and YIplac211-Psm1-Psm3 vectors were integrated into budding yeast.

### Protein expression, purification, labeling and cross-linking

Fission yeast cohesin tetramer complexes including wild type, walker B mutant (Psm1 E1161Q Psm3 E1128Q, denoted as EQ-cohesin), Psm3 acetyl-acceptor site mutant, Rad21 N-terminal truncated mutant (Rad21 amino acids 18-646, denoted N17-cohesin), Kleisin-circle complex, SMC circle complex, Mis4-Ssl3, Mis4-N191(amino acids 192-1587), Pds5 and Wapl were expressed and purified following previously described methods (Chao et al., 2015; Murayama and Uhlmann, 2014, 2015) All fission yeast cohesin complexes for FRET measurement (Head FRET wild type and EQ-cohesin, Kleisin-N FRET wild type, EQ and SG cohesin) were expressed and purified by sequential steps on IgG-sepharose, and heparin columns as described (Murayama and Uhlmann, 2014). The peak fractions from the heparin elution in R buffer (20 mM Tris/HCl, pH 7.5, 0.5 mM TCEP, 10% (v/v) glycerol) containing approximately 600 mM NaCl were concentrated to 500 μl by ultrafiltration. Cohesin was supplemented with 2 μM BG-surface Alexa 647, 1 mM DTT and 0.003% Tween20 and incubated at 25 °C for 1 hour. Now the labeling reaction was supplemented with 4 μM BC-surface Dy547 and incubated at 4 °C for 16 hours to complete the labeling. The labeled cohesin was applied to a Superose 6 10/300 GL gel filtration column that was developed in R buffer containing 200 mM NaCl and 0.003% Tween20. The peak fractions were concentrated to 500 μl by ultrafiltration.

To prepare head-crosslinked cohesin, Head FRET wild type cohesin was expressed and purified by IgG-sepharose chromatography as described above. Once loaded onto the heparin column, R buffer containing 100 mM NaCl and 4 μM SC-Cy5 crosslinker was injected and incubated at 25 °C for 1 hour, resulting mainly in SNAP tag coupling. After this incubation, the column was washed clear of crosslinker and heparin-bound cohesin was eluted and further incubated overnight at 4 °C to allow CLIP tag coupling with SC-Cy5. The peak fractions of heparin purification step were concentrated to 500 μl by ultrafiltration and applied to a Superose 6 10/300 GL gel filtration column that was developed in R buffer containing 200 mM NaCl. The peak fractions were concentrated to 500 μl by ultrafiltration.

### Topological cohesin loading assay

Topological cohesin loading onto DNA was performed in standard reactions (15 μl final volume) as previously described (Murayama and Uhlmann, 2014) with minor modifications. Cohesin (100 nM), Mis4-Ssl3 (100 nM) and pBluescript dsDNA were mixed on ice in reaction buffer (35 mM Tris-HCl pH 7.5, 0.5 mM TCEP, 25 mM NaCl, 1 mM MgCl_2_, 15% (w/v) glycerol and 0.003% (w/v) Tween 20). The reactions were initiated by addition of 0.5 mM ATP and incubated at 32 °C for 120 minutes. The reactions were terminated by addition of 500 μl of ice-chilled Washing buffer A (35 mM Tris-HCl pH 7.5, 0.5 mM TCEP, 750 mM NaCl, 0.35% (w/v) Triton-X100. Anti-Pk antibody bound to protein A conjugated magnetic beads was added to the terminated reactions and rocked at 4 °C overnight. The beads were one time washed with Washing buffer A and three times with Washing buffer B (35 mM Tris-HCl pH 7.5, 0.5 mM TCEP, 500 mM NaCl and 0.1% (w/v) Triton-X100) and once with Washing buffer C (35 mM Tris-HCl pH 7.5, 0.5 mM TCEP, 50 mM NaCl and 0.1% (w/v) Triton-X100). The cohesin-bound DNA was eluted in 15 μl of elution buffer (10 mM Tris-HCl pH 7.5, 1 mM EDTA, 50 mM NaCl, 0.75% SDS and 1 mg/ml protease K) by incubation at 50 °C for 20 minutes. The recovered DNA was separated by 0.8 % agarose gel electrophoresis in TAE buffer and stained with SYBR gold. Gel images were captured using a Typhoon FLA 9500 biomolecular imager and band intensities quantified using Image J.

### Bulk FRET measurement

All fluorescence measurements were carried out at room temperature in reaction buffer (35 mM Tris-HCl pH 7.5, 0.5 mM TCEP, 25 mM NaCl, 1 mM MgCl_2_, 15% (w/v) glycerol and 0.003% (w/v) Tween 20). 40 μl of reaction mixtures containing 10 nM Dy547 and Alexa 647-labeled cohesin, 100 nM Mis4-Ssl3 and 3 nM DNA substrate were mixed and the reaction was started by addition of 0.5 mM ATP. Alternatively, 0.5 mM ADP or 0.5 mM ADP and 0.5 mM BeF_2_, 0.5 mM BeSO_4_ + 10 mM NaF, 0.5 mM AlCl_3_ + 10 mM NaF, or 0.5 mM Na_3_VO_4_ were included instead of ATP. The reactions were incubated at 32 °C for 20 minutes. The samples were applied to a 384-well plate and fluorescence spectra of the cohesin complex were collected on a CLARIOstar high performance plate reader. Samples were excited at 525 nm and emitted light was recorded between 560 - 700 nm in 0.5 nm increments. To evaluate FRET changes caused by cohesin’s conformational changes across different experimental conditions, we report relative FRET efficiency, I_A_/(I_D_ + I_A_), where I_D_ is the donor emission signal intensity at 565 nm resulting from donor excitation at 525 nm and I_A_ is the acceptor emission signal intensity at 665 nm resulting from donor excitation at 525 nm.

### DNA gripping experiments

For DNA gripping analyses, we immobilized DNA on magnetic beads. a 3 kb linear DNA substrate was prepared by PCR amplification with 5’-biotinylated oligonucleotide TH1 and unmodified TH2 using pBluescript dsDNA as the template. The 3 kb DNA loop substrate was made by PCR amplification with a pair of both 5’-biotinylated oligonucleotides TH1 and TH3 using pBluescript dsDNA as the template. Streptavidin conjugated magnetic beads (Invitrogen) were washed with DNA binding buffer DBB (10 mM Tris-HCl pH 7.5, 2 M NaCl, 1 mM EDTA, 0.03% Tween20) and resuspended in 2 volumes of DBB. 100 ng biotin-labelled DNA was mixed with 20 μl beads and incubated at room temperature for 1 hour. Beads were washed 3 times with reaction buffer (35 mM Tris-HCl pH 7.5, 0.5 mM TCEP, 25 mM NaCl, 1 mM MgCl_2_, 15% (w/v) glycerol and 0.003% (w/v) Tween 20) and resuspended in reaction buffer supplemented with 1 mg/ml BSA and 2.5 mU poly-dIdC:dIdC. After 30 minutes incubation, DNA-beads were washed 3 times with reaction buffer. The standard reaction volume was 15 μl, containing 100 nM cohesin, 100 nM Mis4-Ssl3, 100 nM Mis4-N191, 100 nM Pds5 and 100 nM Wapl in reaction buffer. The reaction mixture was added to the DNA beads (containing 3.3 nM dsDNA molecules) on ice. The reactions were started by addition of 0.5 mM ATP, or 0.5 mM ADP and 0.5 mM BeSO_4_ + 10 mM NaF, and incubated at 32 °C for 20 minutes. After the incubation, beads were washed three times with Washing buffer C (35 mM Tris-HCl pH 7.5, 0.5 mM TCEP, 50 mM NaCl and 0.1% (w/v) Triton-X100) or Washing buffer D (35 mM Tris-HCl pH 7.5, 0.5 mM TCEP, 135 mM NaCl and 0.1% (w/v) Triton-X100) and once with Washing buffer C. The beads were divided into two for detection of protein and DNA. Protein samples were eluted with SDS-sample buffer (50 mM Tris-HCl pH 6.8, 2% SDS, 10% Glycerol, 50 mM DTT, 0.02% Bromophenol Blue) and boiled for 5 minutes. DNA sample eluted in buffer containing 3 mM biotin and incubated overnight at room temperature. DNA-bound proteins were separated by SDS-PAGE and analyzed by immunoblotting using the indicated antibodies. The recovered DNA was analyzed by 0.8% agarose gel electrophoresis as described above.

### EM sample preparation of cohesin in the gripping state

For EM sample preparation, we used a 125 bp linear dsDNA substrate that was generated by PCR amplification with a pair of oligonucleotides TH1 and TH5 using pBluescript dsDNA as the template. 200 nM cohesin, 200 nM Mis4-Ssl3, 200 nM 125bp dsDNA were mixed in reaction buffer on the ice. The reaction was started by addition of 0.5 mM ADP and 0.5 mM BeSO_4_ + 10 mM NaF and incubated at 32 °C for 20 minutes. After incubation, an equal volume of 2 x Washing buffer D (35 mM Tris-HCl pH 7.5, 0.5 mM TCEP, 135 mM NaCl and 0.1% (w/v) Triton-X100) was added for further incubation at 4 °C for 10 minutes. The reaction mixture of a total volume of 50 μl was loaded onto 20 - 50% (weight/volume) linear sucrose gradients prepared in EM buffer (20 mM HEPES-KOH pH 7.5, 25 mM NaCl, 0.5 mM TCEP). Centrifugation was in a MLS-50 rotor (Beckman) at 89,513 x *g* for 16 hours at 4 °C. 50 μl fractions were collected from top to bottom and protein and DNA in each fraction were analyzed by SDS-PAGE followed by silver-staining or agarose gel electrophoresis to identify peak fractions containing the cohesion-loader-DNA complex. Sucrose in the peak fractions was removed by passing three times through spin desalting columns before application to EM grids.

### Negative stain EM data acquisition and image processing

A 300-mesh, continuous carbon copper grid (EM Resolutions, C300Cu100) was glow-discharged at 45 mA for 30 seconds. A 4 μl sample was applied and incubated for 1 minute, followed by blotting of excess volume and grid staining in four 50 μl droplets of 2% uranyl acetate for 5, 10, 15, 20 seconds respectively. The grid was subsequently blotted dry. Micrographs were collected at x30,000 nominal magnification (3.45 Å pixel size) with a defocus range of −0.5 to −2.5 μm using a FEI Tecnai LaB6 G2 Spirit electron microscope operated at 120 kV and equipped with a 2K x 2K GATAN UltraScan 1000 CCD camera.

Contrast transfer function parameters were estimated using Gctf v1.06 (Zhang, 2016) and particles were picked semi-automatically with e2boxer in EMAN2 v2.07 (Tang et al., 2007). Subsequent image processing was performed in RELION v3.0.4 (Zivanov et al., 2018). Particles were initially extracted with a box size of 128 pixels and sorted by reference-free 2D classification with CTF-correction using the additional argument --only_flip_phases. To allow visualization of extended Psm1-Psm3 coiled coils, selected cohesin particles were re-extracted with a box size of 192 pixels and processed through one additional round of 2D classification. A reference-free initial 3D model was also created in RELION and used as an input for 3D refinement using particles with the larger box size.

### Cryo-EM data acquisition and image processing

A 400-mesh lacey copper grid with a layer of ultra-thin carbon (Agar Scientific) was glow-discharged at 45 mA for 1 minute. A 4 μl sample was applied to glow-discharged grid and incubated for 2 minutes, followed by blotting of excess volume for 0.5 seconds using a Vitrobot Mark IV (FEI ThermoFisher) operated at room temperature and 100% humidity. To increase particle concentration, two additional 4 μl samples were applied to the grid for 2 minutes each, with 0.5 seconds blotting in between. After a final blot of 3 seconds the grid was plunge-frozen into liquid ethane. High-resolution cryo-EM data were acquired on a FEI Titan Krios electron microscope operated at 300 kV and equipped with Falcon 3EC Direct Electron Detector. Micrographs were collected at x75,000 nominal magnification (1.09 Å pixel size) as 30-frame movies with a total electron dose of 33.8 e^-^/Å^2^ and a defocus range of −2.0 to −4.0 μm. A second dataset was collected with a phase plate using a GATAN K2 Summit direct electron detector operated in counting mode. Micrographs were collected at x130,000 nominal magnification (1.09 Å pixel size) as 40-frame movies with a total electron dose of 49 e^-^/Å^2^ and −0.5 μm defocus.

For the first dataset (no phase-plate), 30-frame movies were corrected for beam-induced movement using 5 x 5 patch alignment with all frames in MotionCor2 (Zheng et al., 2017). Contrast transfer function parameters were estimated on non-dose-weighted micrographs using Gctf v1.06 and particles were picked with crYOLO (Wagner et al., 2019). Subsequent image processing was performed in RELION v3.0.4 and cryoSPARC v2.14.2. Initially 883,184 particles were extracted from 12,085 micrographs in RELION using a box size of 360 pixels. After reference-free 2D classification in cryoSPARC 792,173 cohesin particles were selected and utilized to reconstruct an ab-initio 3D model, which was subsequently used as a starting model for non-uniform refinement. Following 3D refinement in RELION the particle subset was subjected to two rounds of 3D classification using a mask encompassing the cohesin core only. Ultimately 255,148 particles were selected and refined in cryoSPARC using non-uniform refinement followed by local non-uniform refinement of the cohesin core, resulting in a structure at 3.9 Å resolution. The final half-maps were used to produce a density modified map using the Phenix’s tool ResolveCryoEM (Terwilliger et al., 2020). This map showed significant improvements in side chain density and overall interpretability.

For the second dataset (phase-plate), 40-frame movies were corrected for beam-induced movement using 5 x 5 patch alignment using all frames in MotionCor2. Contrast transfer function parameters were estimated on non-dose-weighted micrographs using CTFFIND v4.1.10 (Rohou and Grigorieff, 2015) and particles were picked with crYOLO. Subsequent image processing was performed in RELION v3.0.4 and cryoSPARC v2.14.2. Initially 330,024 binned-by-2 particles were extracted from 5,972 micrographs in RELION using a box size of 276 pixels (2.18 Å/pixel). After reference-free 2D classification in cryoSPARC 227,159 cohesin particles were selected and utilised to reconstruct an ab-initio 3D model, which was subsequently used as a starting model for non-uniform refinement. The particles were re-extracted with a smaller box size of 180 pixels (2.18 Å/pixel) and 3D refined in RELION, which revealed a flexible element connected to the cohesin core. To further characterized this peripheral element, particles were 3D-classified without image alignment using a mask encompassing only the new density region. 80,325 particles were selected and 3D-autorefined in RELION without a mask. To increase the resolution of the flexible element and assess the conformational changes sampled in the cohesion complex, multibody refinement (Nakane et al., 2018) was performed using masks encompassing either the core or the flexible region. The two signal-subtracted particle stacks generated during multibody refinement in RELION were then exported for non-uniform refinement in cryoSPARC. Although the flexible region could not be resolved to subnanometer resolution, a defined, rigid body could be identified, into which a homology model of Psc3 could be unambiguously docked using the Fit-in map command in UCSF Chimera (Pettersen et al., 2004). The homology model was generated with SWISS-MODEL (Waterhouse et al., 2018) and based on PDB entry 6H8Q (Li et al., 2018).

### Model building and validation

SWISS-MODEL was used to obtain homology models for Psm1, Psm3, Rad21 (PDB entries 4UX3 and 1W1W) (Gligoris et al., 2014; Haering et al., 2004), and Mis4, PDB entry 5T8V (Kikuchi et al., 2016). These models were docked into the cryo-EM map using the Fit in Map command in USCF Chimera (Pettersen et al., 2004). These models were refined against the map using Namdinator (Kidmose et al., 2020) and the resulting model was used as a starting point for manual adjustments in Coot (Emsley et al., 2010). The resulting model was then subjected to an iterative process of real space-refinement using Phenix.real_space_refinement (Adams et al., 2010) with geometry and secondary structure restraints followed by manual inspection and adjustments in Coot. Residues 552-583 from Rad21(chain B) and 209-302 from Mis4 (cain D) were docked into the map by rigid-body fitting of the corresponding homology models. The geometries of the atomic model were evaluated by MolProbity (Williams et al., 2018). Cryo-EM data acquisition, 3D reconstruction information and atomic model refinement statistics are summarized in Table S2. Figures were prepared with UCSF Chimera and ChimeraX (Goddard et al., 2018).

### SDA-based protein-protein crosslink mass spectrometry (CLMS) analysis

#### Sample preparation

Protein crosslinking of the cohesin complex was performed in two conditions. An initial state contained all components except nucleotide. The DNA gripping state was achieved by addition of ADP and BeSO_4_ + NaF. All materials (cohesin, loader and DNA) were dialyzed in SDA cross-linking buffer (35 mM HEPES-KOH pH 7.5, 0.5 mM TCEP, 25 mM NaCl, 1 mM MgCl_2_, 15% (w/v) glycerol and 0.003% (w/v) Tween 20) at 4 °C for 3 hours. Cohesin (200 nM), Mis4-Ssl3 (200 nM) and 125 bp dsDNA (200 nM) were mixed on ice in SDA cross-linking buffer. The reaction in each condition was started in the absence of nucleotide or in the presence of 0.5 mM ADP and 0.5 mM BeSO_4_ + 10 mM NaF at 32 °C. After 20 minutes incubation, SDA was added to 50 μg of the cohesin complex at increasing cross-linker weight ratios. (Protein: SDA = 1:1.3, 1:1.9 and 1: 3.8). The diazirine group in SDA was photo-activated using UV irradiation at 365 nm from an ultraviolet crosslinker (Spectrum). Samples were mounted in a 96-well plate, placed on ice at a distance of 5 cm from the UV-A lamp and irradiated for 20 minutes. After UV irradiation, the sample was further incubated on ice for 2 hours to allow further time for NHS crosslinking. The reaction mixtures from the three protein: crosslinker ratios were combined and quenched with 50 mM ammonium bicarbonate. 4 sample volumes of cold acetone were added and incubated at −20 °C for 1 hour. Precipitated proteins were collected by centrifugation and dried in a vacuum concentrator.

#### CLMS sample analysis

Both samples were resolubilized in 100 μl digestion buffer (8M urea in 100 mM ammonium bicarbonate) to an estimated protein concentration of 1 mg/ml. Dissolved protein sample was reduced by addition of 0.5 μl 1M dithiothreitol (DTT) at room temperature for 30 minutes. The free sulfhydryl groups in the sample were then alkylated by adding 3 μl 500 mM iodoacetamide and incubation at room temperature for 20 minutes in the dark. After alkylation, 0.5 μl 1M DTT was added to quench excess of iodoacetamide. Next, protein samples were digested with LysC (at a 50:1 (m/m) protein to protease ratio) at room temperature for four hours. The sample was then diluted with 100 mM ammonium bicarbonate to reach a urea concentration of 1.5 M. Trypsin was added at a 50:1 (m/m) protein to protease ratio to further digest proteins overnight (∼15 hours) at room temperature. Resulting peptides were desalted using C18 StageTips (Rappsilber et al., 2007).

For each sample, resulting peptides were fractionated using size exclusion chromatography in order to enrich for crosslinked peptides (Leitner et al., 2013). Peptides were separated using a Superdex Peptide 3.2/300 column (GE Healthcare) at a flow rate of 10 μl/minute. The mobile phase consisted of 30% (v/v) acetonitrile and 0.1% trifluoroacetic acid. The earliest six peptide-containing fractions (50 μl each) were collected. Solvent was removed using a vacuum concentrator. The fractions were then analysed by LC-MS/MS.

LC-MS/MS analysis was performed using an Orbitrap Fusion Lumos Tribrid mass spectrometer (Thermo Fisher Scientific), connected to an Ultimate 3000 RSLCnano system (Dionex, Thermo Fisher Scientific). Each size exclusion chromatography fraction was resuspended in 1.6% v/v acetonitrile 0.1% v/v formic acid and analysed with replicated LC-MS/MS acquisitions. Peptides were injected onto a 50-centimetre EASY-Spray C18 LC column (Thermo Scientific) that is operated at 50 °C column temperature. Mobile phase A consists of water, 0.1% v/v formic acid and mobile phase B consists of 80% v/v acetonitrile and 0.1% v/v formic acid. Peptides were loaded and separated at a flowrate of 0.3 μl/min. Peptides were separated by applying a gradient ranging from 2% to 45% B over 90 minutes. The gradient was optimized for each fraction. Following the separating gradient, the content of B was ramped to 55% and 95% within 2.5 minutes each. Eluted peptides were ionized by an EASY-Spray source (Thermo Scientific) and introduced directly into the mass spectrometer.

The MS data is acquired in the data-dependent mode with the top-speed option. For each three-second acquisition cycle, the full scan mass spectrum was recorded in the Orbitrap with a resolution of 120,000. The ions with a charge state from 3+ to 7+ were isolated and fragmented using higher-energy collisional dissociation (HCD). For each isolated precursor, one of three collision energy settings (26%, 28% or 30%) was selected for fragmentation using data dependent decision tree based on the m/z and charge of the precursor. The fragmentation spectra were then recorded in the Orbitrap with a resolution of 50,000. Dynamic exclusion was enabled with single repeat count and 60 second exclusion duration.

MS2 peak lists were generated from the raw mass spectrometric data files using the MSConvert module in ProteoWizard (version 3.0.11729). The default parameters were applied, except that Top MS/MS Peaks per 100 Da was set to 20 and the denoising function was enabled. Precursor and fragment m/z values were recalibrated. Identification of crosslinked peptides was carried out using xiSEARCH software (https://www.rappsilberlab.org/software/xisearch) (version 1.7.0) (Mendes et al., 2019). The “initial state” and “gripping state” samples were processed separately. For each sample, peak lists from all LC-MS/MS acquisitions were searched against the sequence and the reversed sequence of cohesin and loader subunits (Psm1, Psm3, Rad21, Psc3, Mis4 and Ssl3). The following parameters were applied for the search: MS accuracy = 4 ppm; MS2 accuracy = 10 ppm; enzyme = trypsin (with full tryptic specificity); allowed number of missed cleavages = 2; missing monoisotopic peak = 2; cross-linker = SDA (the reaction specificity for SDA was assumed to be for lysine, serine, threonine, tyrosine and protein N termini on the NHS ester end and any amino acids for the diazirine end); fixed modifications = carbamidomethylation on cysteine; variable modifications = oxidation on methionine and SDA loop link. Identified crosslinked peptide candidates were filtered using xiFDR (Fischer and Rappsilber, 2017). A false discovery rate of 1% on residue-pair level was applied with “boost between” option selected. A list of identified crosslinked residue pairs is reported in Table S3.

### DNA-protein crosslink mass spectrometry (DPC-MS) analysis

#### Sample preparation

For DNA-protein crosslinking, we prepared two types of dsDNA probes. A 125 bp linear dsDNA was amplified by PCR with 5’-biotinylated oligonucleotide TH1 and non-modified oligonucleotide TH5 using pBluescript dsDNA as the template. PCR reaction mixtures contains 5 ng/ml template DNA, 0.3 μM of each primer, 0.2 mM each of dATP, dCTP and dGTP, 0.02 mM dTTP, 0.18 mM aminoallyl-dUTP and 0.025 unit/μl Go-taq DNA polymerase in 1x Go-taq buffer (Promega). A 3 kb circular dsDNA was prepared by primer extension on single stranded DNA. 5’-biotinylated TH1 oligonucleotide primer was annealed to single strand DNA templates of pBluescript, prepared using M13KO7 helper phage (Murayama et al., 2018). For second strand synthesis, the primer-template mix was incubated in 20 mM Tris-HCl (pH 7.5), 10 mM MgCl_2_, 1 mM DTT, 0.4 mM each of dATP, dCTP, dGTP and aminoallyl-dUTP, 0.1 mg/ml BSA and 0.04 unit/μl T7 DNA polymerase (New England Biolabs) at 37 °C for 3 hours. After synthesis, the buffer of the DNA samples was exchanged to SDAD cross-linking buffer (100 mM NaHCO_3_ pH 8.3) using MicroSpin S400 columns (GE Healthcare). 1 μg of dsDNA was incubated with 2 mM SDAD cross-linker in 25 μl of SDAD cross-linking buffer at 25 °C overnight. The diazirin-decorated dsDNA (SDAD-DNA) probe was dialyzed in DNA dialysis buffer (10 mM Tris-HCl pH 7.5, 0.1 mM EDTA).

For DNA-protein crosslinking 200 nM cohesin, 200 nM Mis4-Ssl3, 20 ng/μl linear 125 bp or circular 3 kb SDAD-DNA probe were mixed in reaction buffer, and DNA gripping reaction was initiated by addition of 0.5 mM ADP and 0.5 mM BeSO_4_ + 10 mM NaF at 32 °C for 30 minutes. An equal volume of 2 x Washing buffer D (35 mM Tris-HCl pH 7.5, 0.5 mM TCEP, 135 mM NaCl and 0.1% (w/v) Triton-X100) was added to the reaction mixture and incubated at 4 °C for 10 minutes. The sample was mounted on a 96-well plate, placed on ice at a distance of 5 cm from the UV-A lamp and irradiated for 10 minutes, as described above for SDA protein-protein crosslinking. After UV irradiation, the buffer of the samples was exchanged with Protease buffer (100 mM ammonium bicarbonate pH 8) using MicroSpin S400 columns. Lys-C protease was added (1:20 (m/m) protease to protein ratio) and incubated at 37 °C overnight. To remove non-crosslinked peptide from the DNA, an equal volume of 2 x RIPA buffer (100 mM Tris-HCl pH 8, 100 mM NaCl, 0.2% SDS) was added to the sample, followed by incubation at 50 °C for 30 minutes. The DNA with crosslinked peptides was purified by Superdex75 size exclusion chromatography developed with 20 mM Tris-HCl pH 7.5, 200 mM NaCl. The recovered DNA-peptide complexes in the void fraction were supplemented with NaCl to 1 M final concentration and 0.1% (w/v) Tween-20. The biotinylated DNA was recovered using streptavidin M280 magnetic beads (Invitrogen) at 25 °C for 1 hour. DNA-beads was washed 3 times with 1 x RIPA buffer and 5 times with peptide elution buffer (20 mM Tris-HCl pH 7.5, 200 mM NaCl). DNA-crosslinked peptides were now eluted by addition of peptide elution buffer containing 25 mM DTT and incubation at 37 °C for 30 minutes.

In the experiment comparing cohesin’s initial binding state and the gripping state, the incubation and crosslinking was performed as above without or with 0.5 mM BeSO_4_ + 10 mM NaF. After the 32 °C incubation, the sample was directly mounted on a 96-well plate without washing buffer addition and the plate was UV irradiated on ice for 10 minutes. The irradiated sample was then treated in the same way as described above.

#### DPC-MS sample analysis

Peptide solutions in the DTT peptide elution buffer were transferred into Total Recovery vials (Waters) for injection without further clean-up or concentration. Samples were analyzed by online nanoflow LC-MS/MS using an Orbitrap Fusion Lumos mass spectrometer (Thermo Scientific) coupled to an Ultimate 3000 RSLCnano (Thermo Scientific). 15 µl of sample was loaded via autosampler into a 20 µl sample loop and pre-concentrated onto an Acclaim PepMap 100 75 µm x 2 cm nanoviper trap column with loading buffer, 2% v/v acetonitrile, 0.05% v/v trifluoroacetic acid, 97.95% water (Optima grade, Fisher Scientific) at a flow rate of 7 µl/min for 6 minutes in the column oven held at 40 °C. Peptides were gradient eluted and separated with a C_18_ 75 µm x 50 cm, 2 µm particle size, 100 Å pore size, reversed phase EASY-Spray analytical column (Thermo Scientific) at a flow rate of 275 nl/min and with the column temperature held at 40 °C, with a spray voltage of 2100 V using the EASY-Spray Source (Thermo Scientific). Gradient elution buffers were A 0.1% v/v formic acid, 5% v/v DMSO, 94.9% v/v water and B 0.1% v/v formic acid, 5% v/v DMSO, 20% v/v water, 74.9% v/v acetonitrile (all Optima grade, Fisher Scientific aside from DMSO, Honeywell Research Chemicals). The gradient elution profile used was 8% B to 40% B over 60 minutes.

The instrument method used an MS1 Orbitrap scan resolution of 120,000 at FWHM m/z 200, quadrupole isolation, mass range 375-1500 m/z, RF Lens 40%, AGC target 4e5, maximum injection time 50 ms and spectra were acquired in profile. Monoisotopic Peak Determination was set to the peptide mode, and only precursors with charge states 2-6 were permitted for selection for fragmentation. Dynamic Exclusion was enabled to exclude after n=1 times for 20 s with high and low ppm mass tolerances of 10 ppm. MS2 scans were acquired in the ion trap following HCD fragmentation with fixed collision energy of 32% and was performed on all selected precursor masses using a cycle time based on data-dependent mode of acquisition set to 3 s. The parameters used for the HCD MS2 scan were quadrupole isolation with an isolation window width of 1.2 m/z, first mass 110 m/z, AGC target 2e3, maximum injection time 300 ms and the scan data was acquired in centroid mode at the rapid scan rate.

A FASTA database containing only the sequences of the six subunits of the cohesin complex and loader was used for the PEAKS search conducted within PEAKS Studio (Bioinformatics Solutions Inc). A modification corresponding to the diazirine moiety after reduction (C_7_H_13_NOS, 159.07178) was created and considered as a variable modification along with the oxidation of methionine residues. Other parameters of the search were digestion enzyme LysC with a maximum of 2 missed cleavages, peptide mass tolerance 5ppm and fragment mass tolerance 0.6 Da.

### SC-Cy5 crosslinking experiments

SC-Cy5 crosslinker was synthesized as previously described (Gautier et al., 2009). Using crosslinkable cohesin complexes (kleisin-circle or SMC-circle) and a DNA-loop substrate, we performed DNA gripping assay as described above. Following the DNA gripping reaction, DNA-beads were washed 3 times with Washing buffer D (35 mM Tris-HCl pH 7.5, 0.5 mM TCEP, 135 mM NaCl and 0.1% (w/v) Triton-X100), and supplemented with 4 μM SC-Cy5 and 1 mM DTT in Washing buffer D. Crosslinking was carried out at 32 °C for 60 minutes. DNA-beads were then divided into three parts. One part was immediately eluted with SDS sample buffer containing 3 mM biotin and served as the input sample. The second sample was washed 5 times with Washing buffer D. The third sample was washed 5 times with SDS buffer (35 mM Tris-HCl pH 7.5, 0.5 mM TCEP, 100 mM NaCl and 0.1% SDS). The second and third samples were then supplemented with SDS sample buffer containing 3 mM biotin and boiled for 10 minutes to elute DNA and protein. Samples were analyzed by SDS-PAGE, followed by in-gel detection of Cy5 or immunoblotting with indicated antibodies.

To further evaluate topological DNA entrapment by the circularized kleisin, beads following SDS washes were divided into two, equilibrated with DNA digestion buffer (35 mM Tris-HCl pH 7.5, 0.5 mM TCEP, 100 mM NaCl, 10 mM MgCl_2_, 0.1 mg/ml BSA, 0.1% Triton X-100) and treated without or with 1 U/μl restriction enzyme PstI in DNA digestion buffer. After a 20 minute incubation at 32 °C, the beads and supernatant fractions were separated, SDS sample buffer containing 3 mM biotin added to each and samples boiled for 10 minutes.

### ATPase assay

Cohesin (150 nM) and Mis4-Ssl3 (100 nM) were mixed with pBluescript dsDNA in reaction buffer (15 μl in final volume). The reactions were initiated by addition of 0.25 mM ATP, spiked with [γ-33P]-ATP, and incubated at 32 °C. Aliquots (2 μl) were taken after 0, 15, 30 and 60 minutes and terminated by addition of 6 μl of 0.5 M EDTA pH 8.0. The products were separated by thin layer chromatography on TCL polyethylenimine cellulose F sheets (Merck), developed with 400 mM LiCl in 1 M formic acid. Plates were analyzed using a Typhoon FLA 9500 Phosphor-imager (GE Healthcare).

---

**Figure S1.**
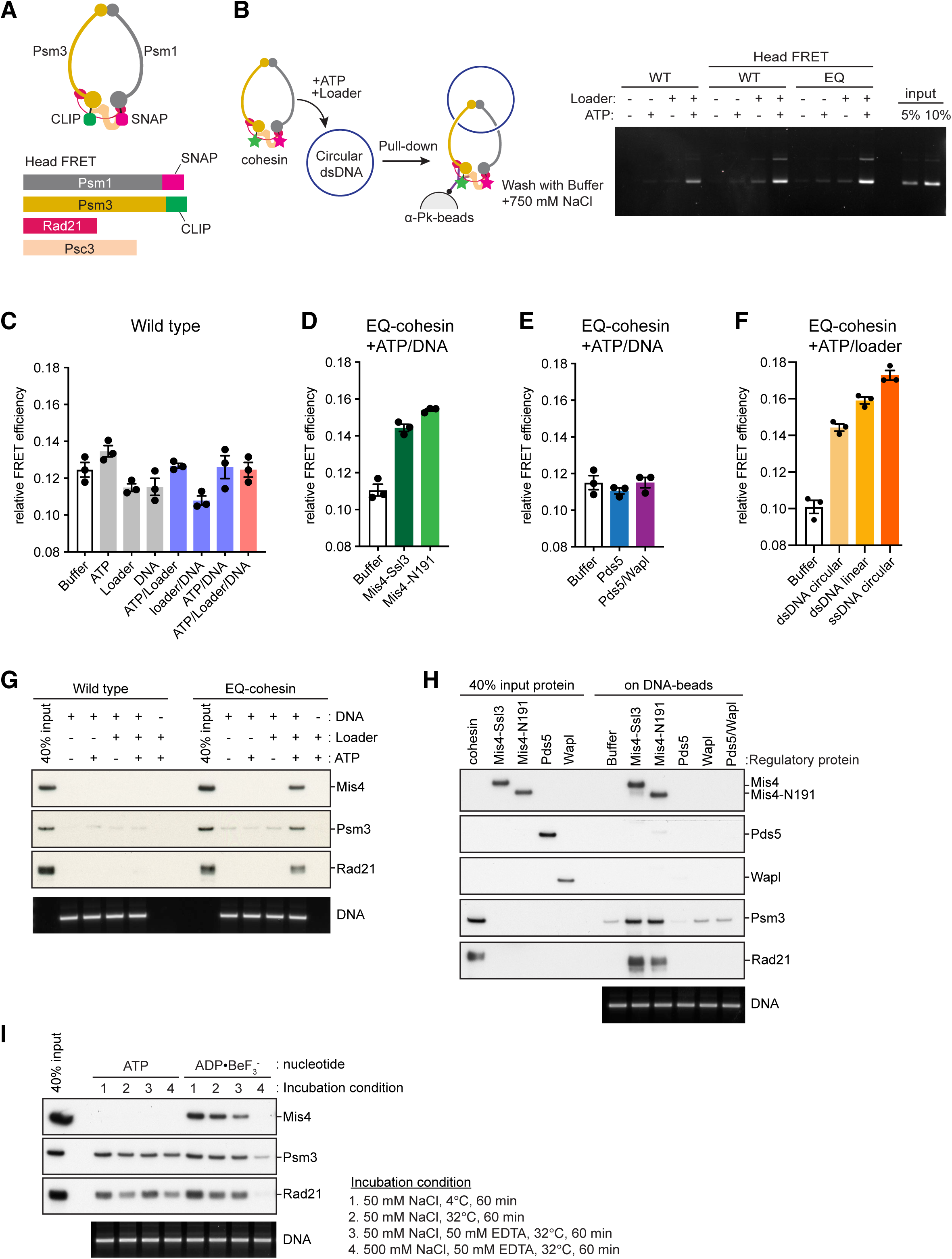
Characterization of Cohesin’s DNA Gripping State (A) Schematic of the modified cohesin complex to record FRET between the ATPase heads. Psm1 and Psm3 were fused to a C-terminal SNAP- and CLIP-tag, respectively. (B) Schematic of a cohesin loading reaction and an experiment to compare the performance of untagged cohesin (WT) with head-labeled wild type (Head FRET, WT) and Walker B mutant cohesin (Head FRET, EQ). Following incubation, cohesin was retrieved by immunoprecipitation, washed with buffer containing 500 mM NaCl, and the retained DNA was analyzed by agarose gel electrophoresis. (C) Head FRET efficiencies of wild type cohesin in absence or presence of combinations of ATP, loader and a 3 kb circular plasmid DNA. (D) Ability of the Mis4-Ssl3 cohesin loader complex, or Mis4-N191, to augment head FRET efficiency of EQ-cohesin in the presence of ATP and a 3 kb circular plasmid DNA. (E) Effect of the cohesin unloading factors Pds5 or Pds5-Wapl on head FRET efficiency of EQ-cohesin in the presence of ATP and a 3 kb circular plasmid DNA. (F) 3 kb circular plasmid dsDNA was compared to linear 3 kb dsDNA and a 3 kb circular ssDNA for its ability to stimulate head FRET efficiency of EQ-cohesin in the presence of ATP and loader. Results from three independent repeats of the experiment, their means and standard deviations are shown in (C) – (F). (G) DNA gripping experiment as in Figure 1D, but comparing wild type and EQ-cohesin in the presence of ATP and additional indicated components. (H) Comparison of the ability of Mis4-Ssl3, Mis4-N191, Pds5 and Pds5-Wapl to promote DNA gripping state formation in an experiment performed with wild type cohesin in the presence of ADP▪BeF_3_^-^. (I) Comparison of cohesin’s DNA binding stability following incubation with loader and either hydrolyzable ATP or non-hydrolyzable ADP▪BeF ^-^ using a bead-bound DNA loop substrate. Following the loading reaction and an initial wash with cold buffer containing 500 mM NaCl, the beads were subjected to a 60 minute incubation under one of the four different indicated washing conditions. Beads were then recovered and bound DNA and protein analyzed as above.

**Figure S2.**
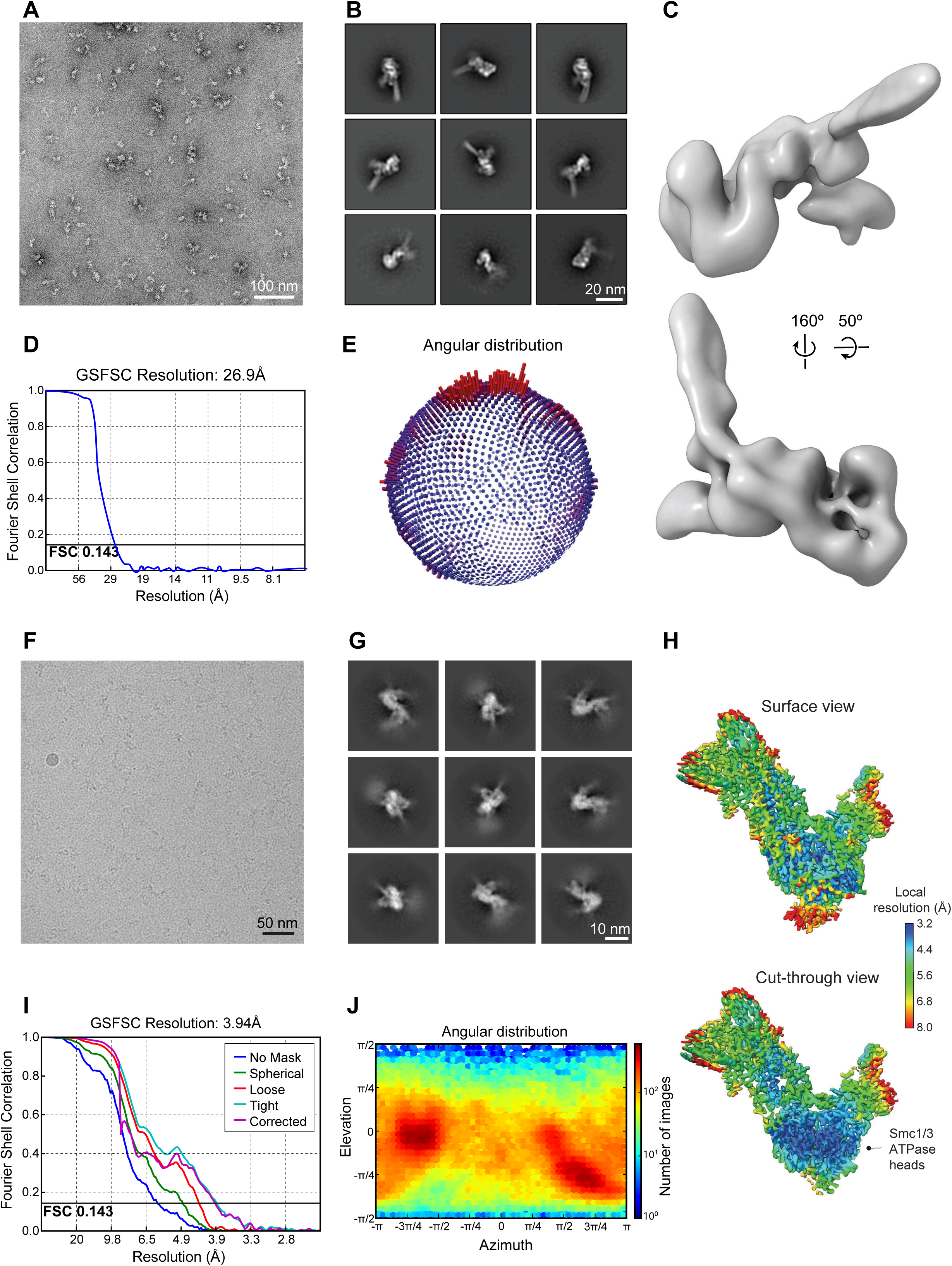
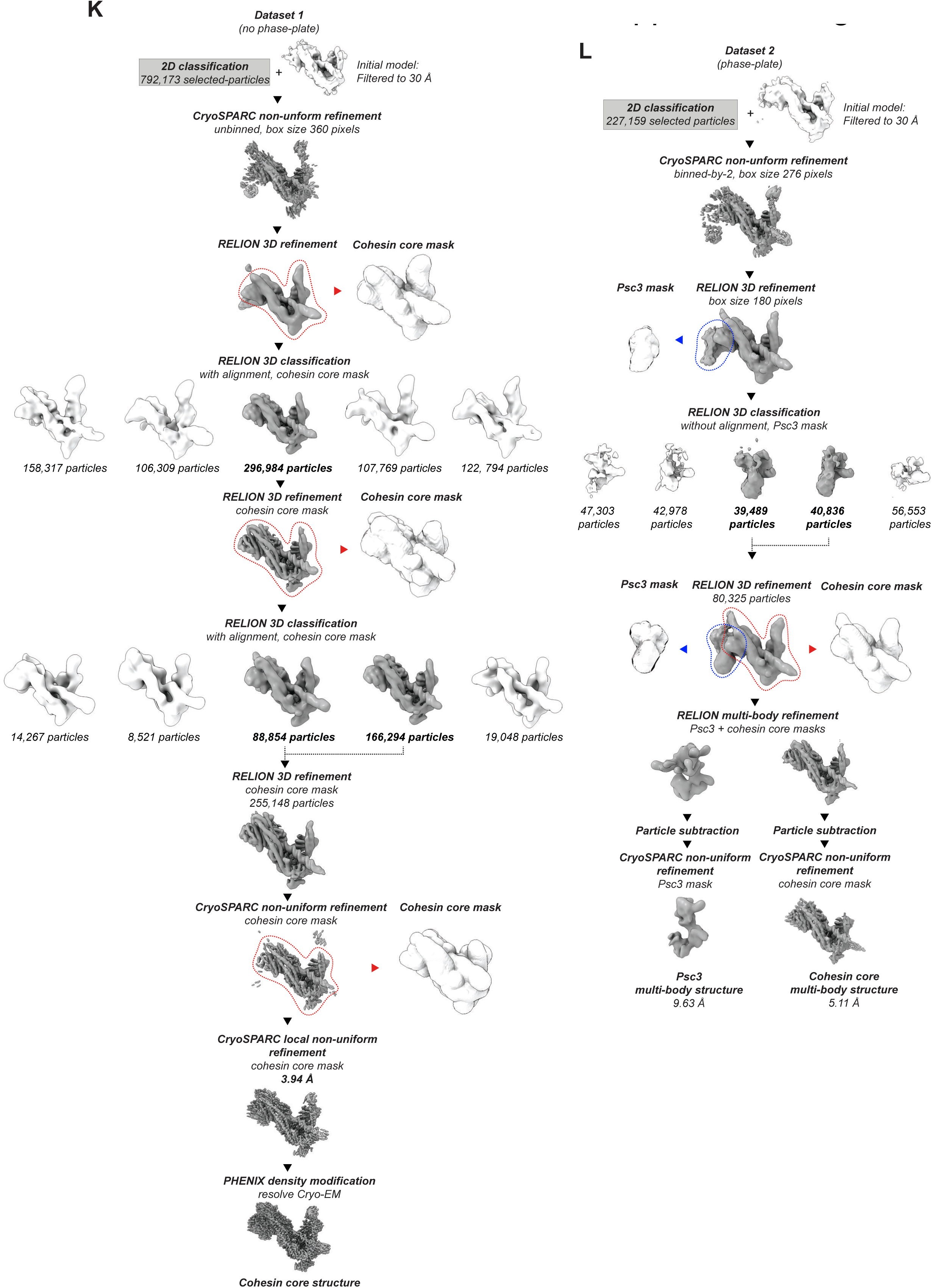
Negative Staining and Cryo-EM approaches to analyze the cohesin-**DNA gripping state** (A) Representative negative stain micrograph. (B) Negative stain 2D class averages. (C) Negative stain 3D reconstruction. (D) Fourier Shell correlation plot with resolution indicated according to the 0.143 criterion. (E) Angular distribution as depicted in RELION. (F) Aligned cryo-EM movie sum obtained from frames collected on a Falcon 3 direct electron detector. (G) Cryo-EM 2D averages. (H) Surface rendering and cut-through view of the cryo-EM cohesin core reconstruction colored according to the local resolution. (I) Fourier Shell correlation plot with resolution indicated according to the 0.143 criterion. The graph was generated using cryoSPARC. (J) Angular distribution. (K) Image processing workflow for the cryo-EM core structure. (L) Multibody refinement workflow, which led to the identification of a separate rigid body identified as Psc3.

**Figure S3.**
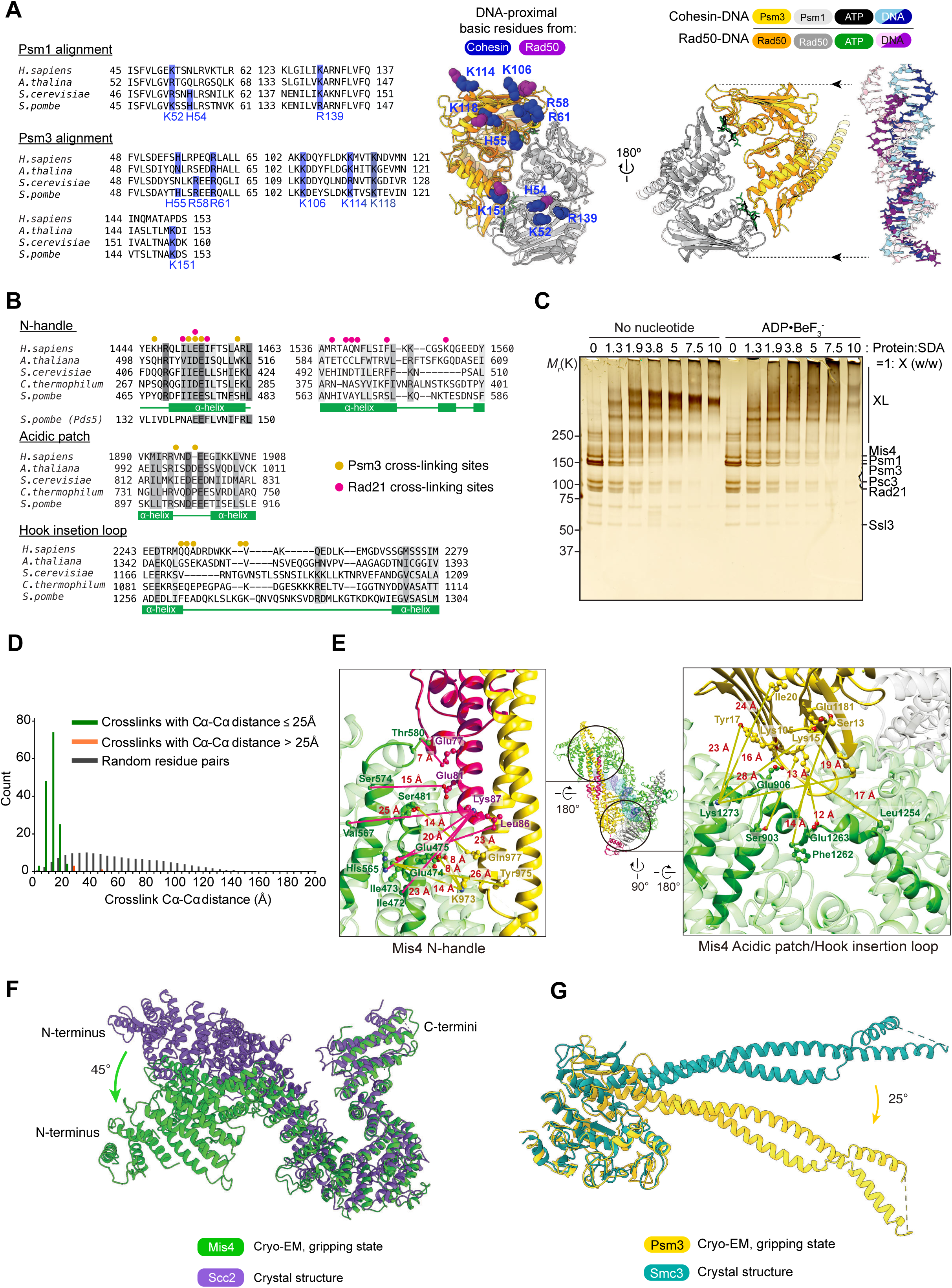
Analyses of the DNA Gripping State (A) On the left, sequence alignment of cohesin SMC heads, highlighting conserved positively charged residues that contribute to DNA contact. On the right, ATPase-head superposition of the nucleotide bound Psm3-Psm1 (gold) and *M. jannashii* Rad50 (orange, PDB: 5F3W) (Liu et al., 2016). DNA-interacting residues are highlighted for Psm3-Psm1 (blue) and Rad50 (purple). Cohesin DNA is cyan and blue, Rad50 DNA is pink and purple. (B) Sequence conservation of three Mis4 elements involved in gripping state formation. Crosslink positions identified in the CLMS experiment are highlighted. A sequence in Pds5, homologous to the Psm3 interacting motif in the Mis4 handle, is also indicated. (C) CLMS samples of the initial binding and DNA gripping state. The SDA crosslinker was added at the indicated ratios and crosslinking induced by UV irradiation at 4 °C for 20 minutes. Samples were analyzed by SDS-PAGE followed by silver staining. Reactions at 1:1.3, 1:1.9 and 1:3.8 ratio were scaled up, combined and used for mass spectrometry analysis. (D) Distribution of Cα-Cα distances of intra-subunit crosslinks measured in the 3D model of the cohesin gripping state. Distances within the theoretical SDA crosslinking limit (25 Å) are colored green, those >25 Å are colored in orange. A normalized distribution of Cα-Cα distances of random residue pairs within these subunits (gray) is shown as reference. Crosslinks from Mis4 are not included in this analysis due to the tendency of this subunit to self-aggregate. (E) CLMS crosslinks between Mis4 and Psm3 (golden lines) and between Mis4 and Rad21 (red lines), mapped on the atomic model of the gripping state. Insets contain magnified views of the Mis4 handle (left) and the Mis4 acidic patch and hook insertion loop (right). (F) Superposition between Mis4 in the gripping state and the orthologous *C. thermophilum* Scc2 in its crystal structure conformation (PDB: 5T8V) (Kikuchi et al., 2016). Superposition of the two structures by the hook domain highlight a reconfiguration of the N-terminal handle. (G) Comparison of Psm3 in the gripping state with *S. cerevisiae* Smc3 in its head-engaged crystal structure conformation (PDB: 4UX3) (Gligoris et al., 2014). Superposition of the two structures by their ATPase head highlight a reconfiguration of the coiled coil neck.

**Figure S4.**
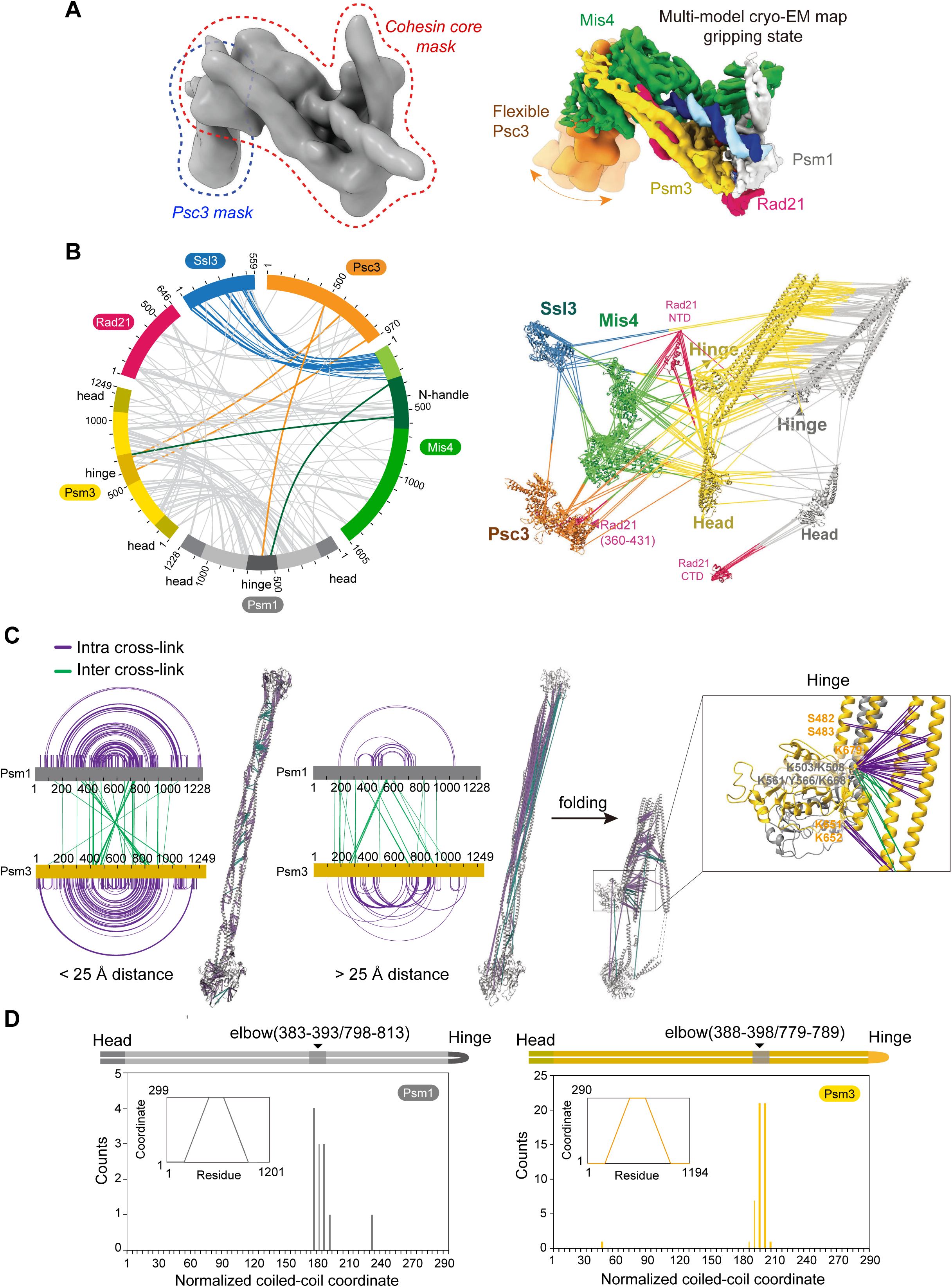
Supporting Analyses of the Hybrid Structural Model (A) Multibody refinement cryo-EM map including Psc3. On the left, 3D structure of the non-masked gripping state core structure. Red and blue traces mark the initial masks used for multibody refinement. On the right, result of the multibody refinement process, including the gripping state core structure and a the flexibly tethered Psc3 density. (B) Circos plot of inter-subunit interactions in the DNA gripping state. Highlighted in blue, crosslinks between Ssl3 and the Mis4 N-terminus; orange, between the Psm1 hinge and Psc3; green, between the Psm3 hinge and the Mis4 handle. The crosslinks are also displayed on an expanded atomic model of the cohesin complex. (C) Diagrammatic and structural representation of crosslinks within (purple) and between (green) Psm1 and Psm3. An extended model of the coiled coils accommodates most crosslinks within 25 Å distance (left), however a series of crosslinks breach this distance constraint (middle). Folding the coiled coil at its predicted elbow allows the latter crosslinks to fall below 25 Å (right). The direction of folding is constrained by crosslinks between the hinge and coiled coil (inset). (D) Identification of the cohesin elbow region from CLMS data. Plot of midpoints of crosslinks with residue pairs that are more than 50 amino acid apart in a normalized coiled coil coordinate system. An inset graph shows the piecewise interpolation function used to map arm residues to the unified coordinate.

**Figure S5.**
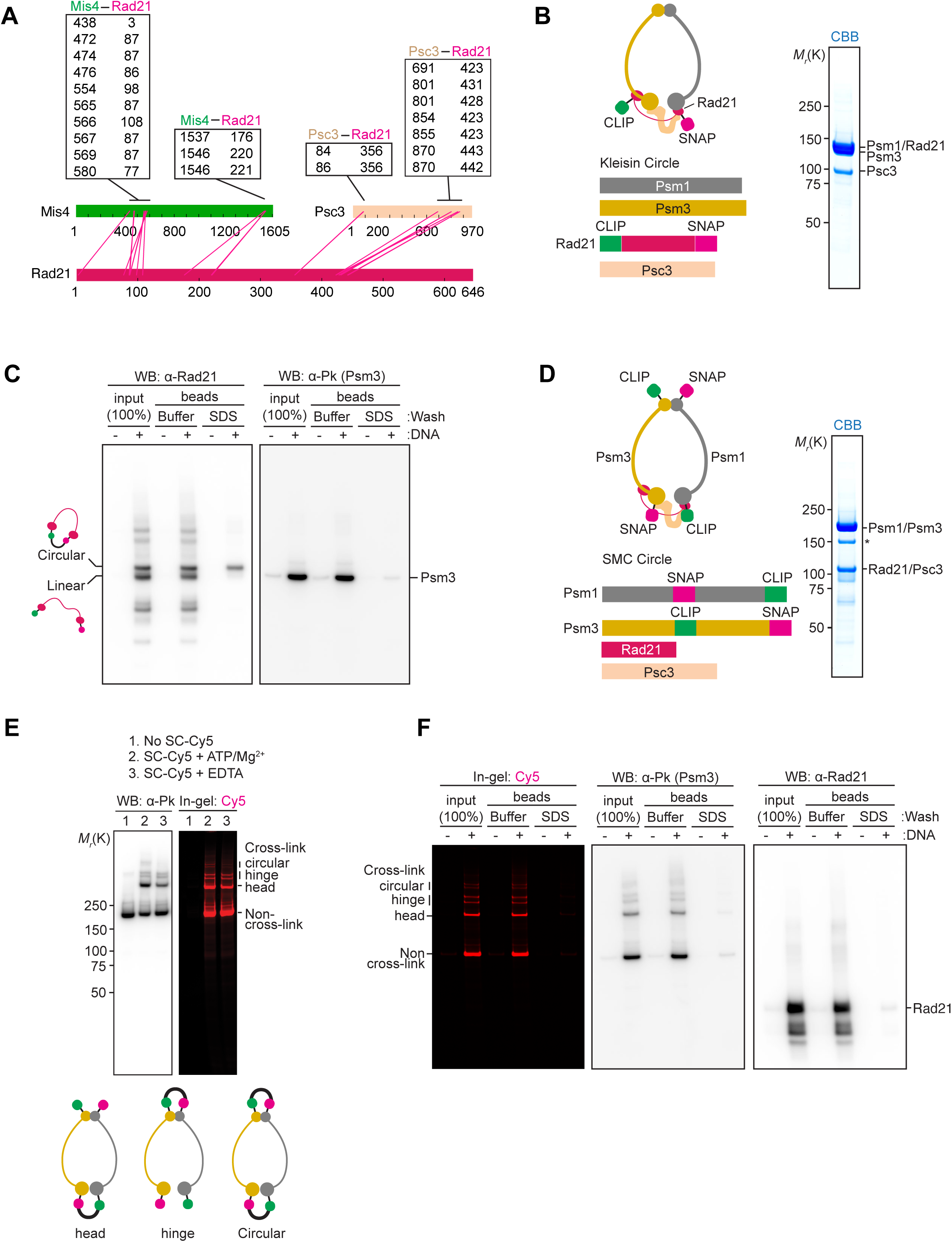
Supporting Information on the Kleisin Path (A) List and graphical depiction of protein crosslinks between Rad21 and Mis4 and Psc3 in the gripping state. (B) Schematic of the cohesin complex containing CLIP and SNAP tags at the kleisin N- and C-terminus, respectively, to circularize the kleisin. The purified complex was analyzed by SDS-PAGE and Coomassie blue (CBB) staining. (C) Immunoblot analysis of the kleisin circularization experiment in Figure 5C. DNA-bound proteins were detected using an antibody against Rad21 or an antibody against the Pk epitope fused to Psm3. Following the SDS wash, Psm3 and linear Rad21 were lost from the DNA beads. (D) Schematic of a cohesin complex that allows SMC circularization. Psm1 has a SNAP-tag inserted between amino acids 593 and 594 at its hinge and a C-terminal CLIP tag. Psm3 in turn contains a CLIP-tag between amino acids 631 and 632 at its hinge as well as a C-terminal SNAP tag. The purified complex was analyzed by SDS-PAGE and Coomassie blue (CBB) staining. (E) Characterization of SMC circularization. SC-Cy5 crosslinking was performed either in the presence of ATP and MgCl_2_ (lane 2) or instead in buffer containing EDTA (lane 3). The latter should prevent head engagement, thus bands of reduced intensity likely correspond to head crosslinks. Crosslinking of both heads and hinge is expected at lower frequency than either single linkage, completing the assignment of the circularized, slowest migrating forms. (F) In gel Cy5 detection and immunoblotting of a DNA gripping experiment using a bead-immobilized DNA loop substrate as in Figure 5D. Unlike what we observed following Rad21 circularization, SMC circularization did not result in SDS resistant protein retention on DNA.

**Figure S6.**
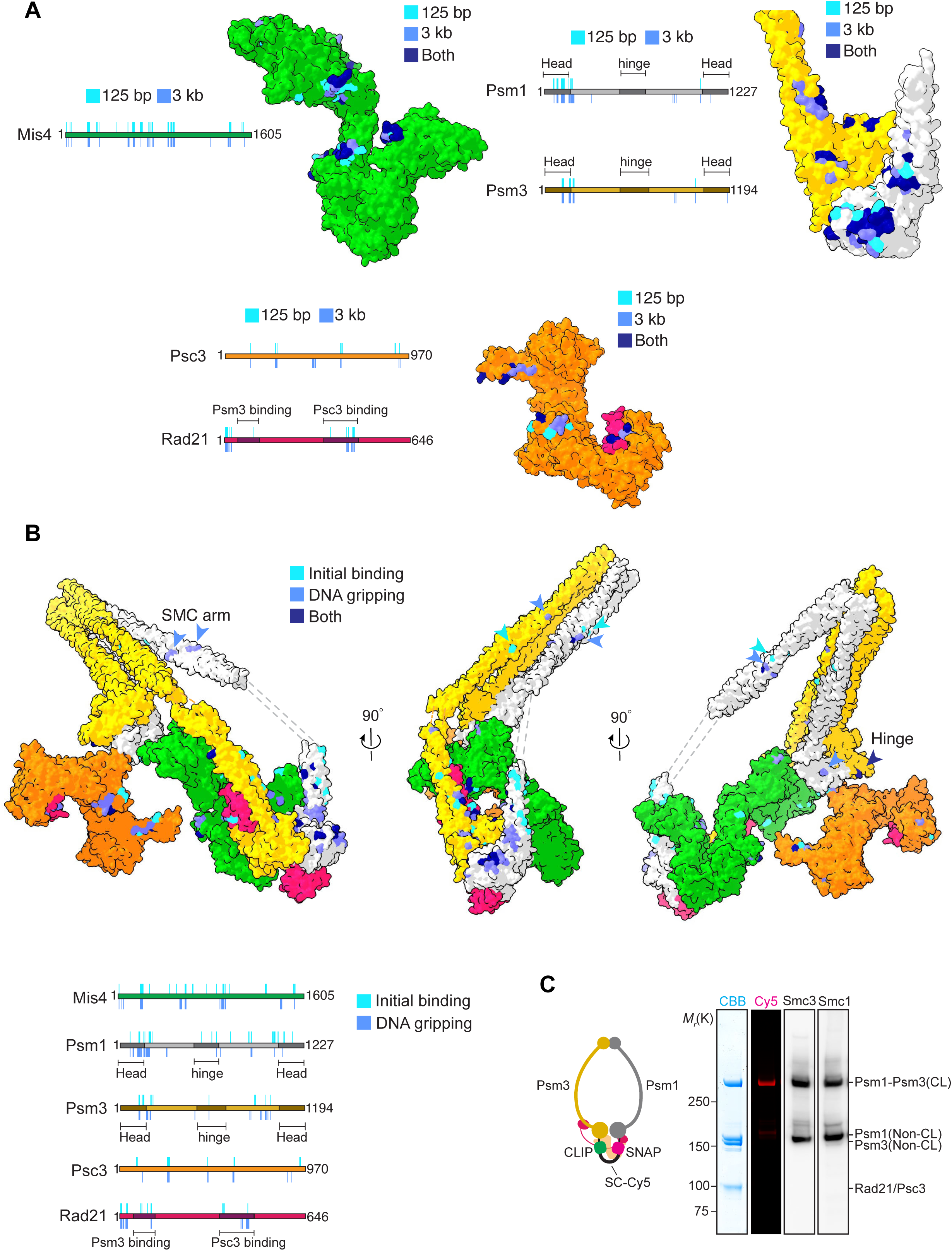
Additional Information on the DPC-MS Results (A) Schematic depiction and surface representation of individual subunits of the DNA crosslinks obtained with either a 125 bp linear DNA or a 3 kb plasmid DNA in the gripping state. (B) Schematic depiction and surface representation on the hybrid model of the DNA crosslinks observed in the initial DNA binding mode and the gripping state. Arrowheads highlight crosslinks along the SMC coiled coils. (C) Schematic of cohesin used for the head crosslinking experiment and analysis of the crosslink products following SC-Cy5 incubation by SDS-PAGE followed by Coomassie blue staining (CBB), in gel Cy5 detection and immunoblotting.

**Figure S7.**
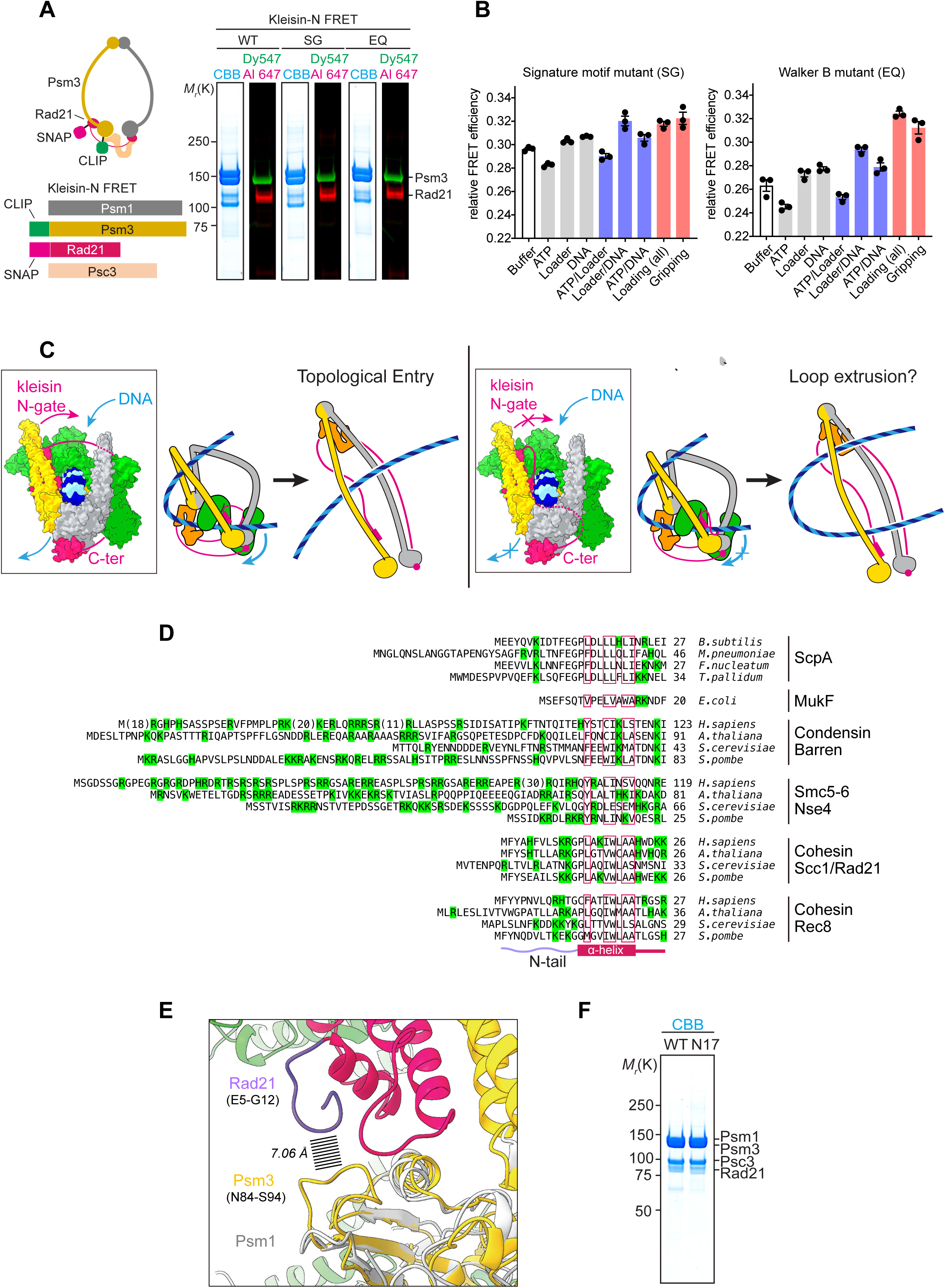
Further Analyses of the Kleisin N-gate and N-tail (A) Schematic of the kleisin N-gate FRET construct. The purified and labeled wild type (WT), signature motif (SG) or Walker B motif mutant (EQ) complexes were analyzed by SDS-PAGE followed by Coomassie blue staining (CBB) or in gel fluorescence detection. (B) As Figure 7A, but FRET efficiencies were recorded using the signature motif and Walker B motif mutant cohesin complexes. Results from three independent repeats of the experiment, their means and standard deviations are shown. (C) Model for the DNA fate when DNA has passed the kleisin N-gate (left), or failed to pass the kleisin N-gate (right), before reaching the ATPase heads. ATP hydrolysis and head gate opening leads to completion of DNA entry into the cohesin ring only in the first scenario. In the second, hypothetical scenario, DNA is unable to enter the cohesin ring and might instead be extruded as a DNA loop. (D) Sequence alignment of kleisin N-tails from various SMC complexes and species. The alignment is centered on the first *<ι>α</i>*-helix (boxed in red). Positively charged amino acids are highlighted green. (E) Analysis of purified wild type (WT) and N17-cohesin by SDS-PAGE followed by Coomassie blue staining (CBB). (F) Atomic model of the Rad21 N-tail in proximity to a Psm3 surface loop. Superposition of the Psm1 head is shown, that lacks a similar loop.

## REFERENCES

1. Adams, P.D., Afonine, P.V., Bunkóczi, G., Chen, V.B., Davis, I.W., Echols, N., Headd, J.J., Hung, L.W., Kapral, G.J., Grosse-Kunstleve, R.W., et al. (2010). PHENIX: a comprehensive Python-based system for macromolecular structure solution. Acta Crystallogr 66, 213–221.

2. Anderson, D.E., Losada, A., Erickson, H.P., and Hirano, T. (2002). Condensin and cohesin display different arm conformations with characteristic hinge angles. J Cell Biol 156, 419–424.

3. Arumugam, P., Gruber, S., Tanaka, K., Haering, C.H., Mechtler, K., and Nasmyth, K. (2003). ATP hydrolysis is required for cohesin’s association with chromosomes. Curr Biol 13, 1941–1953.

4. Beckouët, F., Srinivasan, M., Roig, M.B., Chan, K.-L., Scheinost, J.C., Batty, P., Hu, B., Petela, N., Gligoris, T., Smith, A.C., et al. (2016). Releasing activity disengages cohesin’s Smc3/Scc1 interface in a process blocked by acetylation. Mol Cell 61, 563–574.

5. Ben-Shahar, T.R., Heeger, S., Lehane, C., East, P., Flynn, H., Skehel, M., and Uhlmann, F. (2008). Eco1-dependent cohesin acetylation during establishment of sister chromatid cohesion. Science 321, 563–566.

6. Buheitel, J., and Stemmann, O. (2013). Prophase pathway-dependent removal of cohesin from human chromosomes requires opening of the Smc3-Scc1 gate. EMBO J 32, 666–676.

7. Bürmann, F., Lee, B.G., Than, T., Sinn, L., O’Reilly, F.J., Yatskevich, S., Rappsilber, J., Hu, B., Nasmyth, K., and Löwe, J. (2019). A folded conformation of MukBEF and cohesin. Nat Struct Mol Biol 26, 227–236.

8. Bürmann, F., Shin, H.C., Basquin, J., Soh, Y.M., Giménez-Oya, V., Kim, Y.G., Oh, B.H., and Gruber, S. (2013). An asymmetric SMC–kleisin bridge in prokaryotic condensin. Nat Struct Mol Biol 20, 371–379.

9. Çamdere, G.Ö., Carlborg, K.K., and Koshland, D. (2018). Intermediate step of cohesin’s ATPase cycle allows cohesin to entrap DNA. Proc Natl Acad Sci USA 115, 9732–9737.

10. Chan, K.-L., Roig, M.B., Hu, B., Beckouët, F., Metson, J., and Nasmyth, K. (2012). Cohesin’s DNA exit gate is distinct from its entrance gate and is regulated by acetylation. Cell 150, 961–974.

11. Chao, W.C.H., Murayama, Y., Muñoz, S., Costa, A., Uhlmann, F., and Singleton, M.R. (2015). Structural studies reveal the functional modularity of the Scc2-Scc4 cohesin loader. Cell Rep 12, 719–725.

12. Chao, W.C.H., Murayama, Y., Muñoz, S., Jones, A.W., Wade, B.O., Purkiss, A.G., Hu, X.-W., Borg, A., Snijders, A.P., Uhlmann, F., et al. (2017). Structure of the cohesin loader Scc2. Nat Commun 8, 13952.

13. Davidson, I.F., Bauer, B., Goetz, D., Tang, W., Wutz, G., and Peters, J.-M. (2019). DNA loop extrusion by human cohesin. Science 366, 1338–1345.

14. Eeftens, J.M., Katan, A.J., Kschonsak, M., Hassler, M., de Wilde, L., Dief, E.M., Haering, C.H., and Dekker, C. (2016). Condensin Smc2-Smc4 dimers are flexible and dynamic. Cell Rep 14, 1813–1818.

15. Emsley, P., Lohkamp, B., Scott, W.G., and Cowtan, K. (2010). Features and development of Coot. Acta Crystallogr 66, 486–501.

16. Fischer, L., and Rappsilber, J. (2017). Quirks of error estimation in cross-linking/mass spectrometry. Anal Chem 89, 3829–3833.

17. Furuya, K., Takahashi, K., and Yanagida, M. (1998). Faithful anaphase is ensured by Mis4, a sister chromatid cohesion molecule required in S phase and not destroyed in G_1_ phase. Genes Dev 12, 3408–3418.

18. Gautier, A., Nakata, E., Lukinavicius, G., Tan, K.T., and Johnsson, K. (2009). Selective cross-linking of interacting proteins using self-labeling tags. J Am Chem Soc 131, 17954–17962.

19. Gligoris, T.G., Scheinost, J.C., Bürmann, F., Petela, N., Chan, K.L., Uluocak, P., Beckouët, F., Gruber, S., Nasmyth, K., and Löwe, J. (2014). Closing the cohesin ring: structure and function of its Smc3-kleisin interface. Science 346, 963–967.

20. Goddard, T.D., Huang, C.C., Meng, E.C., Pettersen, E.F., Couch, G.S., Morris, J.H., and Ferrin, T.E. (2018). UCSF ChimeraX: Meeting modern challenges in visualization and analysis. Protein Sci 27, 14–25.

21. Gruber, S., Arumugam, P., Katou, Y., Kuglitsch, D., Helmhart, W., Shirahige, K., and Nasmyth, K. (2006). Evidence that loading of cohesin onto chromosomes involves opening of its SMC hinge. Cell 127, 523–537.

22. Guacci, V., Chatterjee, F., Robison, B., and Koshland, D.E. (2019). Communication between distinct subunit interfaces of the cohesin complex promotes its topological entrapment of DNA. Elife 8, e46347.

23. Haering, C.H., Farcas, A.M., Arumugam, P., Metson, J., and Nasmyth, K. (2008). The cohesin ring concatenates sister DNA molecules. Nature 454, 297–301.

24. Haering, C.H., Löwe, J., Hochwagen, A., and Nasmyth, K. (2002). Molecular architecture of SMC proteins and the yeast cohesin complex. Mol Cell 9, 773–788.

25. Haering, C.H., Schoffnegger, D., Nishino, T., Helmhart, W., Nasmyth, K., and Löwe, J. (2004). Structure and stability of cohesin’s Smc1-kleisin interaction. Mol Cell 15, 951–964.

26. Hara, K., Zheng, G., Qu, Q., Liu, H., Ouyang, Z., Chen, Z., Tomchick, D.R., and Yu, H. (2014). Structure of cohesin subcomplex pinpoints direct shugoshin-Wapl antagonism in centromeric cohesion. Nat Struct Mol Biol 21, 864–870.

27. Hassler, M., Shaltiel, I.A., Kschonsak, M., Simon, B., Merkel, F., Thärichen, L., Bailey, H.J., Macošek, J., Bravo, S., Metz, J., et al. (2019). Structural basis of an asymmetric condensin ATPase cycle. Mol Cell 74, 1175–1188.

28. Hinshaw, S.M., Makrantoni, V., Kerr, A., Marston, A.L., and Harrison, S.C. (2015). Structural evidence for Scc4-dependent localization of cohesin loading eLIFE *4*, e06057.

29. Hirano, T. (2016). Condensin-based chromosome organization from bacteria to vertebrates. Cell 164, 847–857.

30. Hopfner, K.-P., Karcher, A., Shin, D.S., Craig, L., Arthur, L.M., Carney, J.P., and Tainer, J.A. (2000). Structural Biology of Rad50 ATPase: ATP-driven conformational control in DNA double-strand break repair and the ABC-ATPase superfamily. Cell 101, 789–800.

31. Huis in ‘t Veld, P.J., Herzog, F., Ladurner, R., Davidson, I.F., Piric, S., Kreidl, E., Bhaskara, V., Aebersold, R., and Peters, J.-M. (2014). Characterization of a DNA exit gate in the human cohesin ring. Science 346, 968–972.

32. Jeppsson, K., Kanno, T., Shirahige, K., and Sjögren, C. (2014). The maintenance of chromosome structure: positioning and functioning of SMC complexes. Nat Rev Mol Cell Biol 15, 601–614.

33. Kidmose, R.T., Juhl, J., Nissen, P., Boesen, T., Karlsen, J.L., and Panyella, B.P. (2020). Namdinator - Automatic Molecular Dynamics flexible fitting of structural models into cryo-EM and crystallography experimental maps. bioRxiv.

34. Kikuchi, S., Borek, D.M., Otwinowski, Z., Tomchick, D.R., and Yu, H. (2016). Crystal structure of the cohesin loader Scc2 and insight into cohesinopathy. Proc Natl Acad Sci USA 113, 12444–12449.

35. Kim, Y., Shi, Z., Zhang, H., Finkelstein, I.J., and Yu, H. (2019). Human cohesin compacts DNA by loop extrusion. Science 366, 1345–1349.

36. Lee, B.-G., Roig, M.B., Jansma, M., Petela, N., Metson, J., Nasmyth, K., and Löwe, J. (2016). Crystal structure of the cohesin gatekeeper Pds5 and in complex with kleisin Scc1. Cell Rep 14, 1–8.

37. Leitner, A., Walzheoeni, T., and Aebersold, R. (2013). Lysine-specific chemical cross-linking of protein complexes and identification of cross-linking sites using LC-MS/MS and the xQuest/xProphet software pipeline. Nat Protoc 9, 120–137.

38. Li, Y., Muir, K.W., Bowler, M.W., Metz, J., Haering, C.H., and Panne, D. (2018). Structural basis for Scc3-dependent cohesin recruitment fo chromatin. eLIFE 7, e38356.

39. Liu, Y., Sung, S., Kim, Y., Li, F., Gwon, G., Jo, A., Kim, A.K., Kim, T., Song, O.K., Lee, S.E., et al. (2016). ATP-dependent DNA binding, unwinding, and resection by the Mre11/Rad50 complex. EMBO J 35, 743–758.

40. Mendes, M.L., Fischer, L., Chen, Z.A., Barbon, M., O’Reilly, F.J., Giese, S.H., Bohlke-Schneider, M., Belsom, A., Dau, T., Combe, C.W., et al. (2019). An integrated workflow for crosslinking mass spectrometry. Mol Syst Biol 15, e8994.

41. Minamino, M., Higashi, T.L., Bouchoux, C., and Uhlmann, F. (2018). Topological in vitro loading of the budding yeast cohesin ring onto DNA. Life Sci Alliance 1, e201800143.

42. Muir, K.W., Li, Y., Weis, F., and Panne, D. (2020). The structure of the cohesin ATPase elucidates the mechanism of SMC-kleisin ring opening. Nat Struct Mol Biol 27, 233–239.

43. Muñoz, S., Minamino, M., Casas-Delucchi, C.S., Patel, H., and Uhlmann, F. (2019). A role of chromatin remodeling in cohesin loading onto chromosomes. Mol Cell 74, 664–673.

44. Murayama, Y., Samora, C.P., Kurokawa, Y., Iwasaki, H., and Uhlmann, F. (2018). Establishment of DNA-DNA interactions by the cohesin ring. Cell 172, 465–477.

45. Murayama, Y., and Uhlmann, F. (2014). Biochemical reconstitution of topological DNA binding by the cohesin ring. Nature 505, 367–371.

46. Murayama, Y., and Uhlmann, F. (2015). DNA entry into and exit out of the cohesin ring by an interlocking gate mechanism. Cell 163, 1628–1640.

47. Nakane, T., Kimanius, D., Lindahl, E., and Scheres, S.H. (2018). Characterisation of molecular motions in cryo-EM single-particle data by multi-body refinement in RELION. Elife 7, e36861.

48. Ouyang, Z., Zheng, G., Tomchick, D.R., Luo, X., and Yu, H. (2016). Structural basis and IP6 requirement for Pds5-dependent cohesin dynamics. Mol Cell 62, 248–259.

49. Petela, N.J., Gligoris, T.G., Metson, J., Lee, B.G., Voulgaris, M., Hu, B., Kikuchi, S., Chapard, C., Chen, W., Rajendra, E., et al. (2018). Scc2 is a potent activator of cohesin’s ATPase that promotes loading by binding Scc1 without Pds5. Mol Cell 70, 1134–1148.

50. Pettersen, E.F., Goddard, T.D., Huang, C.C., Couch, G.S., Greenblatt, D.M., Meng, E.C., and Ferrin, T.E. (2004). UCSF Chimera--a visualization system for exploratory research and analysis. J Comput Chem 25, 1605–1612.

51. Rappsilber, J., Mann, M., and Ishihama, Y. (2007). Protocol for micro-purification, enrichment, pre-fractionation and storage of peptides for proteomics using StageTips. Nat Protoc 2, 1896–1906.

52. Rohou, A., and Grigorieff, N. (2015). CTFFIND4: Fast and accurate defocus estimation from electron micrographs. J Struct Biol 192, 216–221.

53. Schüler, H., and Sjögren, C. (2016). DNA binding to SMC ATPases - trapped for release. EMBO J 35, 703–705.

54. Seifert, F.U., Lammens, K., Stoehr, G., Kessler, B., and Hopfner, K.-P. (2016). Structural mechanism of ATP-dependent DNA binding and DNA end bridging by eukaryotic Rad50. EMBO J 35, 759–772.

55. Tanaka, K., Hao, Z., Kai, M., and Okayama, H. (2001). Establishment and maintenance of sister chromatid cohesion in fission yeast by a unique mechanism. EMBO J 20, 5779–5790.

56. Tang, G., Peng, L., Baldwin, P.R., Mann, D.S., Jiang, W., Rees, I., and Ludtke, S.J. (2007). EMAN2: an extensible image processing suite for electron microscopy. J Struct Biol 157, 38–46.

57. Terwilliger, T.C., Ludtke, S.J., Read, R.J., Adams, P.D., and Afonine, P.V. (2020). Improvement of cryo-EM maps by density modification. bioRxiv.

58. Tóth, A., Ciosk, R., Uhlmann, F., Galova, M., Schleiffer, A., and Nasmyth, K. (1999). Yeast Cohesin complex requires a conserved protein, Eco1p (Ctf7), to establish cohesion between sister chromatids during DNA replication. Genes Dev 13, 320-333.

59. Uhlmann, F. (2016). SMC complexes, from DNA to chromosomes. Nat Rev Mol Cell Biol 17, 399–412.

60. Ünal, E., Heidinger-Pauli, J.M., Kim, W., Guacci, V., Onn, I., Gygi, S.P., and Koshland, D.E. (2008). A molecular determinant for the establishment of sister chromatid cohesion. Science 321, 566–569.

61. Wagner, T., Merino, F., Stabrin, M., Moriya, T., Antoni, C., Apelbaum, A., Hagel, P., Sitsel, O., Raisch, T., Prumbaum, D., et al. (2019). SPHIRE-crYOLO is a fast and accurate fully automated particle picker for cryo-EM. Commun Biol 2, 218.

62. Waterhouse, A., Bertoni, M., Bienert, S., Studer, G., Tauriello, G., Gumienny, R., Heer, F.T., de Beer, T.A.P., Rempfer, C., Bordoli, L., et al. (2018). SWISS-MODEL: homology modelling of protein structures and complexes. Nucl Acids Res 46, W296-W303.

63. Weitzer, S., Lehane, C., and Uhlmann, F. (2003). A model for ATP hydrolysis-dependent binding of cohesin to DNA. Curr Biol 13, 1930–1940.

64. Williams, C.J., Headd, J.J., Moriarty, N.W., Prisant, M.G., Videau, L.L., Deis, L.N., Verma, V., Keedy, D.A., Hintze, B.J., Chen, V.B., et al. (2018). MolProbity: More and better reference data for improved all-atom structure validation. Protein Sci 27, 293–315.

65. Zhang, J., Shi, X., Li, Y., Kim, B.-J., Jia, J., Huang, Z., Yang, T., Fu, X., Jung, S.Y., Wang, Y., et al. (2008). Acetylation of Smc3 by Eco1 is required for S phase sister chromatid cohesion in both human and yeast. Mol Cell 31, 143–151.

66. Zhang, K. (2016). Gctf: Real-time CTF determination and correction. J Struct Biol 193, 1–12.

67. Zheng, S.Q., Palovcak, E., Armache, J.P., Verba, K.A., Cheng, Y., and Agard, D.A. (2017). MotionCor2: anisotropic correction of beam-induced motion for improved cryo-electron microscopy. Nat Methods 14, 331–332.

68. Zivanov, J., Nakane, T., Forsberg, B.O., Kimanius, D., Hagen, W.J., Lindahl, E., and Scheres, S.H. (2018). New tools for automated high-resolution cryo-EM structure determination in RELION-3. Elife 7, e42166.

